# Genomic clustering by geography not species in taxonomically complex British and Irish eyebrights (*Euphrasia*)

**DOI:** 10.1101/2023.03.19.533315

**Authors:** Yanqian Ding, Chris Metherell, Wu Huang, Peter M. Hollingsworth, Alex D. Twyford

## Abstract

Genomic studies of incipient speciation are fundamental to understand the origin and establishment of species. However, a wide range of evolutionary processes and complex evolutionary interactions remain to be explored outside of genetically tractable evolutionary and ecological model systems. Here, we study taxonomically complex British and Irish eyebrights (*Euphrasia*), as a test case for how different evolutionary factors influence species boundaries across geographic space. *Euphrasia* is a plant genus that has remarkable diversity in ploidy, mating system and ecology. There are 21 British and Irish *Euphrasia* species, but with species that are exceptionally difficult to identify based on morphology or DNA barcoding. Here, we test the hypothesis that species boundaries are highly permeable, and taxa experience extensive gene flow despite potential barriers such as ploidy and contrasting mating systems. To understand geographic genetic structure and the nature of species differences, we applied genotyping-by-sequencing (GBS) and spatial-aware clustering methods to 378 population samples from 18 British and Irish species. We find the selfing heathland specialist *E. micrantha* demonstrates genome-wide divergence in Northern Scotland, indicative of a distinct post-glacial colonisation history and the role of a highly selfing mating system in divergence. In contrast, all other genetic clusters correspond to geographic regions, with extensive gene flow between species and a complete absence of species-specific SNPs. Our results reveal the highly permeable species boundaries present in a recently diverging group, with an overriding signal of geographic genetic structure over and above genetic clustering by species.

## Introduction

Species demonstrate a continuum of genomic divergence, from young species with shallow divergence, to older species characterised by genome-wide differentiation (Coyne & Orr, 2004). Studying young species–those at the earliest stage of the speciation continuum–is now a widely recognised way to understand the genetic underpinnings of the origin and establishment of new species (Andrew & Rieseberg, 2013; Choi et al., 2020; Marques et al., 2016). Many such studies, applying genomic sequencing data to ecological and evolutionary model systems, have now shown incipient species are characterised by genomic differentiation that is limited to few regions of the genome that underlie species differences (barrier loci, and linked loci), with the rest of the genome experiencing extensive homogenising gene flow. This leads to a dominant genome-wide signal (i.e., in neutral loci) related to geography, with individuals clustering by sampling location, rather than by taxonomic affinity. For example, cichlid fish of the genus *Astatotilapia* in Tanzania have genome-wide sequence data that cluster individuals by crater lake, with extremely low genome-wide divergence (F_ST_ = 0.038) with fewer than ~1 % of genomic intervals divergent between benthic-littoral ecomorphs (Malinsky et al., 2015). Similarly in the yellow monkey flower Mimulus guttatus, genome-wide genetic clustering corresponds to geographic locations across western North America (southern, coastal or northern regions), with a large chromosomal inversion, the main genomic region of divergence between morphologically distinct annual and perennial ecotypes (Twyford & Friedman, 2015).

A common feature of these recent genomic speciation studies is a focus on tractable systems where a known ecological variable is likely to drive divergence. Here, studies typically select evolutionary systems that are amenable to study, with attributes such as a small diploid genome size and experimental tractability. Most often these include a pair of taxa with well characterised ecological divergence, for example open vs deep water sticklebacks (McKinnon & Rundle, 2002), or monkeyflowers restricted to seasonally dry vs permanently moist soils (Lowry et al., 2008). While much has been learnt from such groups, this overlooks a large proportion of biodiversity that is less tractable, due to their high species diversity, complexity in terms of genomic composition (e.g., large genomes, polyploids) or for experimental work (e.g., hard to culture, long generation times), or poorly characterised ecological selection pressures. Studies in these diverse but largely overlooked groups will allow us to investigate a wider range of evolutionary processes and investigate more complex evolutionary interactions.

A particular challenge for studying the genomics of species differences are taxonomically complex groups, where species are difficult to classify into discrete species. These species often possess ambiguous, continuous, or cryptic trait differences, and are frequently characterised by hybridisation, polyploidy, self-fertilisation, and the occurrence of complex sexual systems. Familiar examples include water fleas (*Daphnia*) (Marková et al., 2007) and dandelions (*Taraxacum*) (Mártonfiová, 2015), both of which are diverse genera characterised by species complexes. While potentially challenging, taxonomically complex groups can also provide important evolutionary insights. For example, many of the difficulties in species identification are due to recent divergence and ongoing homogenising gene flow, both of which are recognised as benefits of studying the earliest stage of the speciation continuum, where it may be possible to identify regions underlying speciation. Moreover, many other factors common to these groups, such as polyploidy and the transition to self-fertilisation, are known to promote divergence and speciation, and depending on the group, these can be studied independently or in concert. Finally, an appreciable number of species fall in these groups, including many recently evolved narrow endemics, where genetic data may prove useful in aiding targeted conservation measures (González-Pérez & Caujapé-Castells, 2022; Ennos et al., 2005; French et al., 2008)

The plant genus *Euphrasia* (Orobanchaceae) in Britain and Ireland are a renowned example of a taxonomically complex group. These species, commonly known as eyebrights, are small in stature and defined by a complex suite of morphological characters. However, from an evolutionary perspective, many of the underlying evolutionary processes making species hard to identify make them valuable to study. Polyploidy is common throughout the genus, and there are 5 native diploids and 16 allotetraploids in Britain and Ireland, with whole genome sequencing revealing British diploids are one parent of British allotetraploids (Becher et al., 2020; Yeo, 1956). Species readily hybridise, with 71 hybrid combinations identified to date, based on morphology (Metherell & Fred, 2018). Phylogenomic analyses have revealed extremely close relationships between species, with British samples showing 99.8% pairwise sequence identity based on complete plastid genomes (Garrett et al., 2022). British species vary in their mating system, from small flowered selfers through to large flowered mixed maters and outcrossers (French et al., 2004; Yeo, 1966). While their species boundaries and speciation histories are currently poorly understood, currently defined taxa are usually ecologically distinct and found in contrasting environments, for example with species restricted to dry heathland, northern coastal saltmarshes, and poor montane soils. Moreover, these species are generalist hemiparasitic plants that are green and photosynthesise but also derive nutrients from a wide range of host, and as such ecological speciation coupled with other major evolutionary transitions (such as to self-fertilisation) are likely to underlie the origin of new species (Becher et al., 2020; Brown et al., 2020).

Here, we investigate population genomic variation across species of taxonomically complex British and Irish eyebrights. We perform extensive population sampling and genomic sequencing with Genotyping by Sequencing (GBS) (Elshire et al., 2011). Our overarching goal is to investigate the genomic distinctiveness of recently divergent species, and to identify how different evolutionary processes interact to shape genomic variation and maintain species barriers. We first aimed to test the role of polyploidy in the origin and maintenance of genomic variation. Previous studies in *Euphrasia* have shown clustering by ploidy using amplified fragment length polymorphism (AFLP) markers, suggesting ploidy is a major barrier to gene flow (French et al., 2008). However, the close relationship between diploid and the tetraploid sub-genome, coupled with the known occurrence of putative cross-ploidy hybrids and even putative cross-ploidy hybrid species, suggests this may not be absolute (Becher et al., 2020). Second, we aim to identify whether any species are genomically distinct across their range, which would be indicative of reproductive isolation with other incipiently diverging taxa, and if so to link this to potential causal factors such as a selfing mating system. Finally, we look at genomic variation across all populations to test if there are regional geographic genetic clusters across species, indicative of a model of divergence with gene flow (i.e., ongoing hybridisation). Combined, these results provide a rare test of the genomic distinctiveness of species in a taxonomically complex plant group and help identify the evolutionary pressures leading to speciation.

## Materials and Methods

### Sampling and genotyping by sequencing

Our sampling strategy aimed to collect the range of British and Irish *Euphrasia* species, with multiple geographically spaced population samples for the most widespread species. In total, 862 *Euphrasia* individuals from 378 populations were collected, 5 populations of which were collected from outside of Britain and Ireland (Jersey and Belgium), and 3 outgroups (diploids, *E. picta* from Austria, *E. pectinata* and *E. regelli* from China) were included. This collection included extensive sampling of 18 native British and Irish *Euphrasia* species (14 tetraploid species and 4 diploid species) and 51 putative hybrid combinations (44 putative tetraploid hybrids, 2 putative diploid hybrids, 5 putative interploidy hybrids) (Figure 1 & Supplemental Table 1). Samples were collected in the field or sent by botanical recorders as part of the Eye for Eyebrights public engagement project. Leaves or whole plants were sampled into silica gel. Voucher specimens for field collections are available in the herbarium at the Royal Botanic Garden Edinburgh (E). All specimens were reidentified by the *Euphrasia* referee, Chris Metherell.

**Figure 1.**
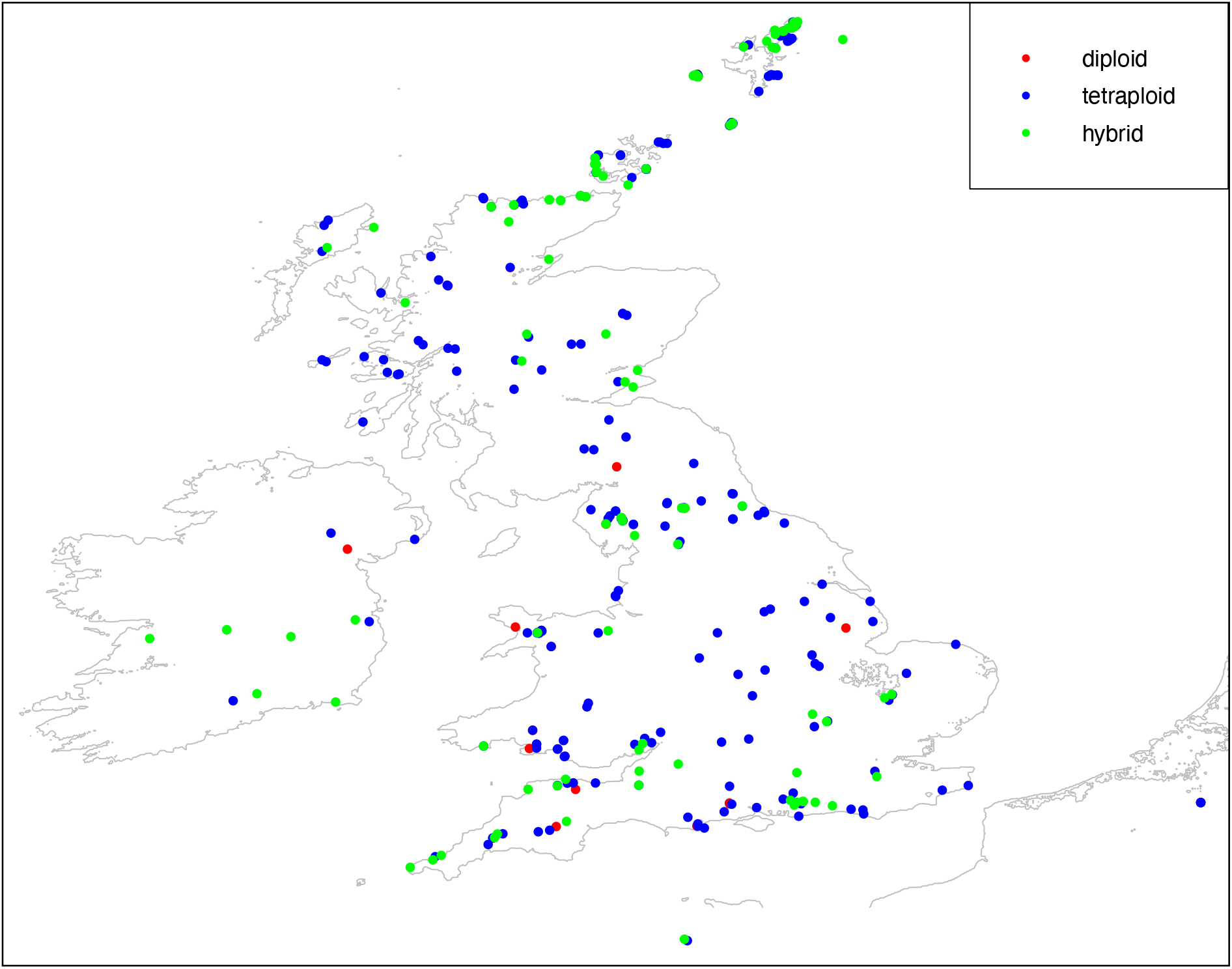
Map of all *Euphrasia* sampling locations. Diploids are limited in their distribution to southern regions, including England, Wales, and Ireland, while tetraploids are widely distributed across Britain and Ireland.

We extracted genomic DNA using the Qiagen DNeasy Plant minikit, prior to sequencing using Genotyping by Sequencing. We followed the protocol of Elshire *et al*. (2011), with individual 95 DNA samples and a water control sample digested with the enzyme ApeKI, barcoded, and pooled as 96 samples per lane of sequencing. Samples were sequenced at Edinburgh Genomics on 11 lanes of either Illumina HiSeq 2500 or HiSeq 4000, using 100bp single end reads. Due to low sequencing coverage, 148 samples were sequenced again, with data combined between technical replicates.

Co-dominant single nucleotide polymorphism (SNP) markers were called using two different draft genomes. We used the tetraploid genome of *E. arctica* from Becher *et al*. (2020); and assembled a new draft genome for the 600Mb diploid *E. anglica* based on 250-bp paired Illumina data presented in Becher *et al*. (2020). This new diploid assembly used DISCOVAR *de novo*, and we assessed the quality of the assembly using BUSCO scores (Seppey et al., 2019). Preliminary SNP calls using sequencing reads from tetraploid samples mapped to the diploid genome resulted in high read mapping rates, and very high heterozygosity (Twyford, Unpublished Results). This is unexpected in (partly) selfing species and is indicative of erroneous SNP calls where short reads from both sub-genomes in the tetraploid incorrectly map to the diploid genome. Therefore, only diploid samples were mapped to the diploid genome (the original DDA dataset herein, i.e., <u>D</u>iploid samples mapped to the <u>D</u>iploid reference genome using <u>A</u>ll scaffolds). To perform comparative analyses between both diploid and tetraploid samples, all samples were mapped to the reference genome of tetraploid *E. arctica*. To avoid biases with missing data in diploids, we extracted the 3,454 sequence scaffolds with similar coverage across diploid and tetraploid individuals identified by Becher *et al*. (2020), which are proposed to represent a conserved sub-genome across taxa and are suitable for comparative analyses. This dataset is called the original ATC dataset herein (i.e., <u>A</u>ll samples, including diploids and tetraploids, mapped to the <u>T</u>etraploid genome, where we extracted the <u>C</u>onserved scaffolds). To understand whether potential biases may have arisen due to selecting this subset of scaffolds, we also looked at TTA dataset (<u>T</u>etraploid samples mapped to the <u>T</u>etraploid genome with <u>A</u>ll tetraploid scaffolds) (Supplemental Text 1).

All mapping was performed using BWA (Li & Durbin, 2009), and SNPs were called with the TASSEL-GBS pipeline (Glaubitz et al., 2014). GBS datasets often have highly heterogeneous patterns of missing data per individual and site. We filtered individuals containing more than 90% missing data and sites containing more than 90% missing data using PLINK v1.9 (Chang et al., 2015; Purcell et al., 2007). This initial filtered dataset was further filtered for fastStructure analyses, removing sites in tight linkage disequilibrium (unlinked SNPs dataset) with the parameter of ‘--indep 50 5 1’, removing SNPs in strong LD within a 50 SNP window, prior to moving 5 SNPs along. To test whether genetic clusters observed in the population genetic structure analyses were affected by extensive sampling of particular geographic regions or species groups (especially the bias of 800 tetraploid samples vs 80 diploids), we down sampled the ATC dataset to a similar number of individuals between the most distinctive genetic clusters recovered in preliminary analyses, before repeating analyses on this dataset (Supplemental Text 2).

### Quantifying genomic differentiation

We characterised whether any eyebright species are defined by diagnostic SNPs using the species-specific allele analysis pipeline (https://github.com/Hazelhuangup/Species_specific_alleles_analysis). This identifies two types of SNPs, (1) those fixed differences between a target species all other taxa (species specific SNPs), (2) major allele frequency differences where a SNP state occurs at >87.5% in the focal species and <12.5% in other species (i.e., a more relaxed threshold allowing for factors such as rare introgression or sequencing/SNP calling errors). The analysis was performed on each dataset, using the sequence alignment and species names as an input, with the analysis repeated for every species.

To quantify the amount of genetic variation partitioned by ploidy and/or taxa, we first performed analysis of molecular variance (AMOVA) on all samples, and on all species (i.e., excluding hybrids). Then in a second analysis we chose three widely distributed and closely related tetraploid species: *E. arctica*, *E. confusa* and *E. nemorosa*, to test the variation among geographic regions and sample populations. Geographical regions were arbitrarily defined to 5 areas of similar size, each containing multiple populations from at least two of these three species (Supplemental Figure 1). AMOVA was then performed with the R package poppr (v2.9.3) (Kamvar et al., 2014, 2015).

### Population structure

To test between genetic clustering of individuals by ploidy level, species, and geography, we performed a range of analyses testing population genetic structure, including principal component analysis (PCA) (Jombart & Ahmed, 2011; Jombart & Bateman, 2008), discriminant analysis of principal components (DAPC) (Jombart et al., 2010), and Bayesian clustering (fastStructure) (Raj et al., 2014). Analyses were performed separately for the diploid-only DDA dataset, and the combined diploid and tetraploid ATC dataset. PCA was performed using glPca() in adegenet (v2.1.5) (Jombart & Ahmed, 2011) and the R package ggplot2 (v3.3.6) (Wickham, 2016) was used for visualisation. fastStructure analyses were run testing genetic clusters (K) from 1 to 10 for the DDA and 1 to 15 for the ATC dataset and choosing model complexity (best K) using the chooseK.py script. Admixture proportions and geographic maps were visualised using the make.structure.plot function in the R packages conStruct (v1.0.5) (Bradburd, 2019) and marmap (v1.0.6) (Pante & Simon-Bouhet, 2013), respectively.

We further tested whether population genetic variation is driven by geography, by investigating the strength of isolation by distance (IBD) across taxa and between genetic clusters identified with fastStructure, above. To evaluate IBD, the R packages geodist (v0.0.7) (Padgham & Sumner, 2021) and hierfstat (v0.5.10) (Goudet & Jombart, 2021) were used to calculate pairwise geographic distances and pairwise F_ST_ for all populations of diploid samples and widely distributed tetraploid samples based on the DDA & ATC unlinked-SNP datasets, respectively. To test whether the genetic clusters identified with fastStructure are robust to potential IBD, we reanalysed our SNPdata with ConStruct (Bradburd et al., 2018). This approach can infer continuous and discrete population genetic structure across sampling space in the presence of IBD (Twyford et al., 2020). Analyses were performed both as nonspatial (without incorporating sampling location information) and spatial (with sampling information) for the DDA dataset and ATC dataset, respectively.

## Results

The newly assembled draft genome for diploid *E. anglica* based on 77-fold coverage short read data was comprised of relatively short contigs (N50 4.2Kb), however the high completeness (94.1% complete BUSCO genes, of which 83.1% were single copy and 11% duplicated) makes it suitable for our application of mapping population-level short GBS reads. Overall, our genotyping-by-sequencing generated a total of 679,126,630 short sequence reads. SNP calling using the TASSEL-GBS pipeline resulted in a total of 100,903 and 27,513 SNPs from the unfiltered DDA (<u>D</u>iploid samples mapped to the <u>D</u>iploid reference genome using <u>A</u>ll scaffolds) and ATC datasets (<u>A</u>ll samples, including diploids and tetraploids, mapped to the <u>T</u>etraploid genome, where we extracted the <u>C</u>onserved scaffolds), respectively. After sites and individuals were filtered for missing data, there were 26,278 and 19,666 SNPs, for the DDA and ATC datasets, respectively. The average missing data per site was 11% (DDA) and 19% (ATC). After individual filtering, 768 individuals from 356 populations remained for downstream analyses. Filtering unlinked SNPs for the fastStructure analyses resulted in 6,016 SNPs for the DDA dataset and 5,220 SNPs for the ATC dataset (Supplemental Table 1).

### Broad scale genetic structure and the role of polyploidy

To explore broad scale patterns of genetic structure, including the influence of polyploidy and the clustering of genetic variation by species, we performed AMOVA, PCA, DAPC and fastStructure analyses. AMOVA showed ploidy only explained 10.6% of genetic variance across species (Table 1). The PCA revealed clustering of individuals by ploidy only in PC3 (9.66% variation, Supplemental Figure 2). Instead of ploidy, the main axis of divergence across all samples was between the selfing taxon *E. micrantha* in northern Britain (particularly individuals found in Shetland and Orkney), as well as its hybrids with *E. scottica* and some other northern individuals of *E. arctica*, *E. fharaidensis*, *E. ostenfeldii*, *E. frigida*, *E. confusa*, and all other samples (PC1, 21.9% variation Figure 2A, Supplemental Figure 3). The fastStructure analyses give similar results to the PCA, and showed the main genetic clusters recovered at K=2 is the northern selfing species, and all other taxa (Figure 2C, 2D, 2E). Northern *E. micrantha* remains genetically cohesive and distinct from other species even at high K-values such as K=9, with clear genomic divergence (F_ST_ = 0.28 between northern *E. micrantha* vs others, Supplemental Figure 4). All 12 sample populations in the north of Scotland are assigned to this distinct cluster, though some populations within Scotland are assigned to other genetic clusters, suggesting the geographic demarcation of these clusters is not clear-cut.

**Table 1.** AMOVA analyses between ploidy and taxa for British and Irish *Euphrasia* using the ATC dataset (19,666 SNPs). We report values for species only / species and hybrids.

|  | df † | % Total variance |
| --- | --- | --- |
| Between ploidy levels | 1/1 | 10.6*/10.5* |
| Between taxa within ploidy | 15/44 | 12.4*/16.9* |
| Within taxa | 223/543 | 76.9*/72.6* |

† df: degrees of freedom.

**Figure 2.**
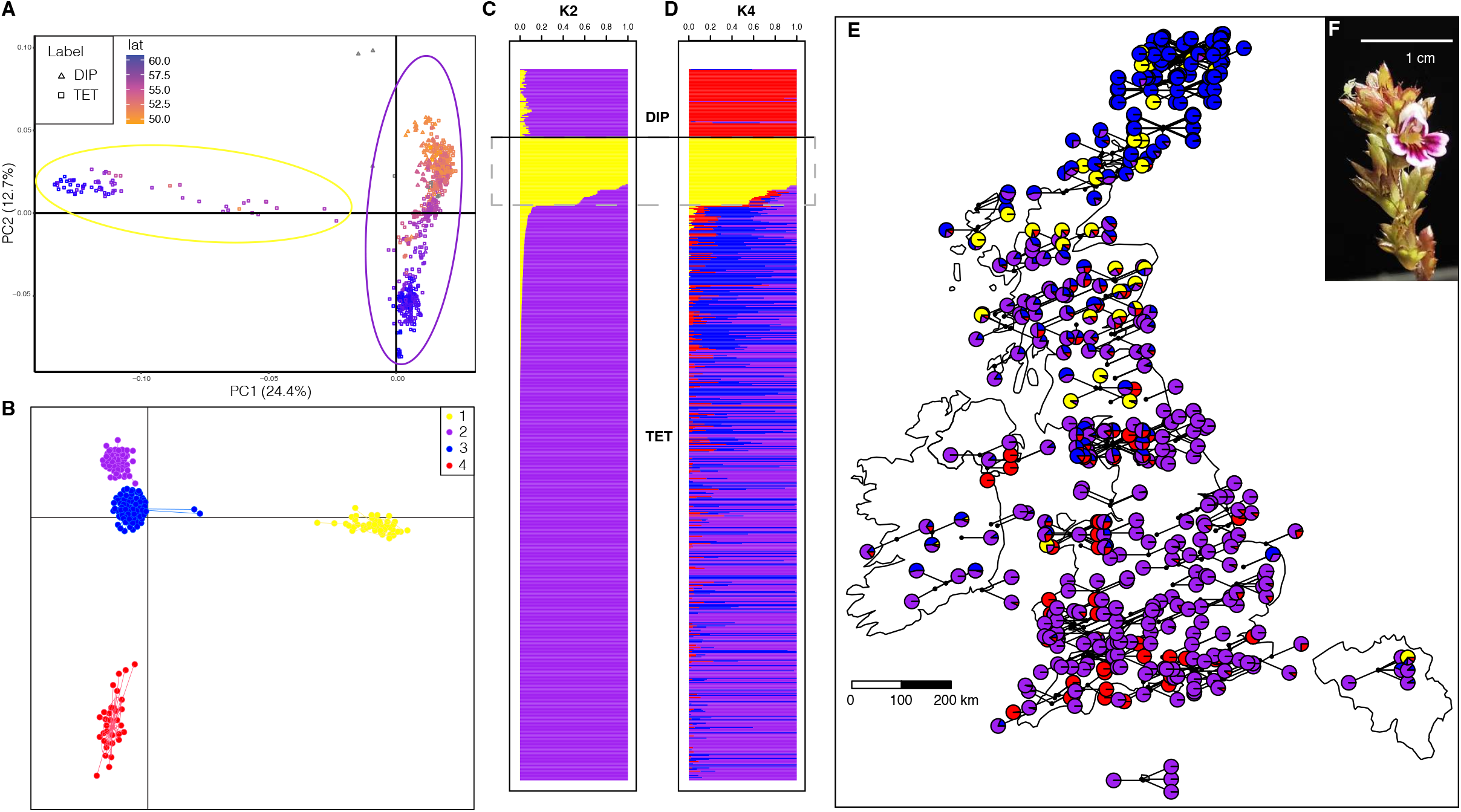
Geographic genetic structure across British and Irish *Euphrasia* sequenced with genotyping by sequencing (GBS). **A)** Principal components analysis (PCA) showing PC1 & 2. Triangles represent diploid samples and rectangles are tetraploid samples. Colour indicates latitude. **B)** Discriminant analysis of principal components (DAPC). Individuals are coloured based on groupings with fastStructure with K=4 (D). **C & D**). fastStructure result for K = 2 and 4. **E).** Genetic cluster map for fastStructure analysis with K=4; pie charts show the proportion of each cluster within a population. **F).** Photograph Of *E. micrantha*. Genetic clusters are coloured consistently across panels, with yellow representing the distinct cluster of northern *E. micrantha*, red are diploid samples.

### Fine scale genetic structure and local geographic clustering

To investigate the drivers of genetic structure, we further explored clustering at higher k-values in the fastStructure analyses. We found that there was little support for the genetic clustering of individuals within species except northern *E. micrantha*, with both the diploids (DDA dataset) and the widely distributed tetraploids (ATC dataset) showing genetic clustering of individuals by geography rather than by species (Figures 3 & 4). In diploids, the optimum K was between 2 to 4, with the analysis with K=4 revealing three genetic clusters corresponding to local geographic clustering in England and one cluster further in northern Scotland and Belgium. There is also a putative cross-ploidy hybrid between *E. arctica* x *E. rostkoviana* clustering with individuals of *E. rostkoviana* sampled from Belgium. In widely distributed tetraploids, the optimum K was between 2 to 7, with similar patterns of individuals clustering by geographic location, except the distinctive cluster of Northern *E. micrantha*. In no cases was there evidence for genetic clusters corresponding to species apart from Northern *E. micrantha*.

**Figure 3.**
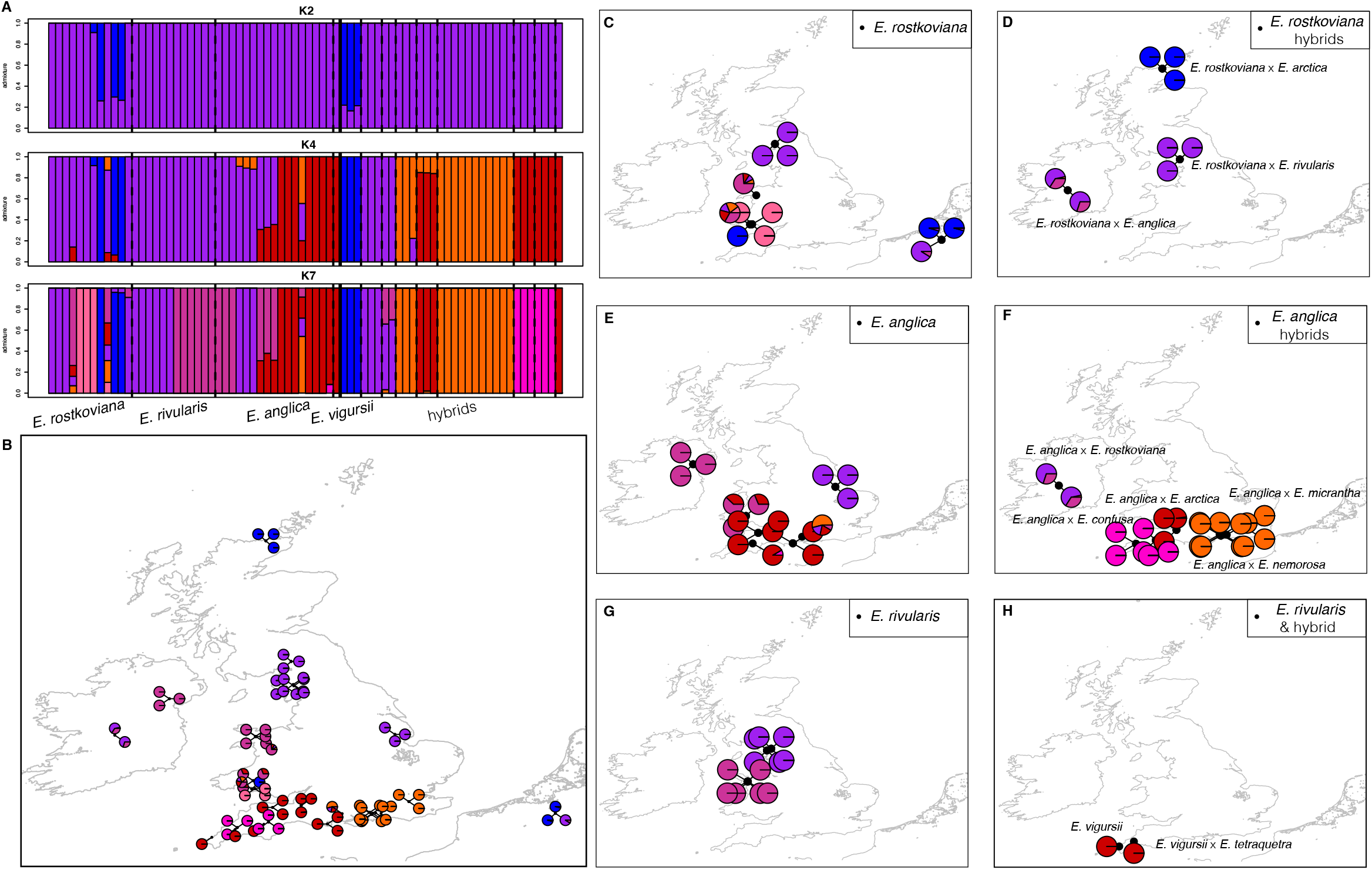
Genetic structure of diploid *Euphrasia* samples**. A)** fastStructure plots of K=2, 4 and 7. **B)** fastStructure map of all diploid samples when K=7. **C-H)** fastStructure map of each species and putative hybrids when K=7.

**Figure 4.**
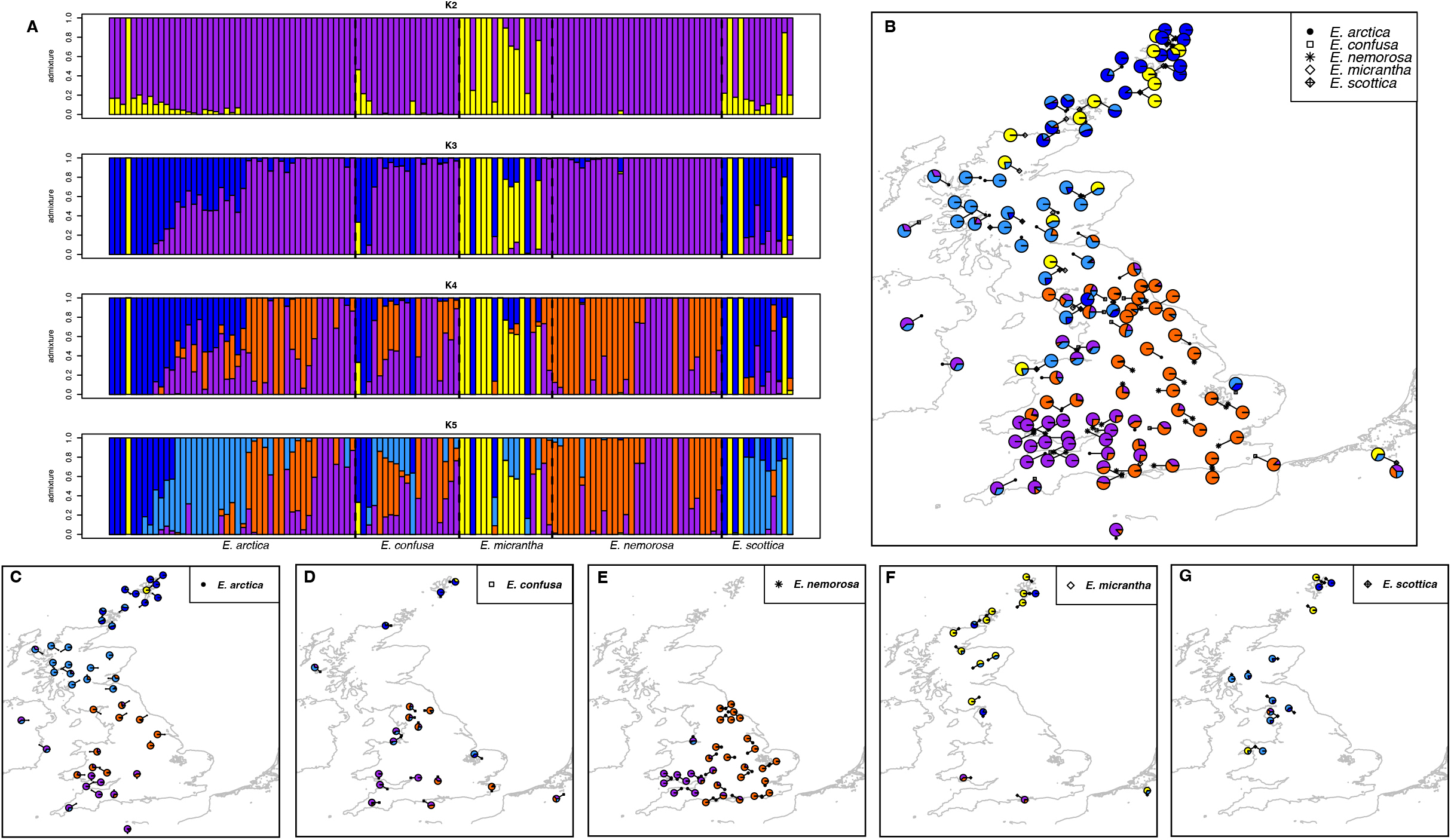
Genetic structure of 5 widely distributed tetraploid species: *E. arctica*, *E. confusa*, *E. nemorosa*, *E. micrantha* and *E. scottica*. **A)** fastStructure plots of K=2-5. **B)** fastStructure map of all 5 tetraploid samples when K=5. **C-G)** fastStructure map of each species when K=5.

We subsequently performed isolation by distance (IBD) analyses of all individuals within a ploidy level, with the aim of testing whether populations follow a simple model of geographic genetic isolation. We found a moderate amount of the genetic variance was explained when all diploid species were analysed in a single combined analysis (r=0.1947, p = 0.004, Supplemental Figure 5). When accounting for the geographic locations of samples in spatially explicit conStruct analyses, we still found no signal of genetic clustering corresponding to any other widespread species (Supplemental Figure 6). However, in tetraploids, excluding the northern selfing genetic cluster primarily composed of *E. micrantha*, the samples showed significant IBD but with very low effect size (r=0.06, p < 0.001, Supplemental Figure 7).

Our test of species-specific SNPs detected only a single diagnostic site, for *E. anglica*, with no other species having species specific SNPs. To quantify the relative partitioning of genetic variance between species and geographic regions, we performed AMOVA on three widespread and closely related tetraploid taxa. AMOVA showed no variation to be partitioned by species, and instead a very high proportion of variation was attributable to differences among populations (54.4%; P < 0.05, Table 2) and among individuals (36%; P < 0.05, Table 2). Within species, 13.5% of the variance was attributable to geographical region (Table 2; P < 0.05).

**Table 2.** Partitioning of SNP variation in *E. arctica*, *E. nemorosa*, *E. confusa* based on an AMOVA considering geographical regions and species.

|  | df | % Total variance |
| --- | --- | --- |
| Between species | 2 | -3.9% |
| Between regions within species | 8 | 13.5* |
| Between populations within region | 23 | 54.4* |
| Within populations | 21 | 36.0* |

## Discussion

The close evolutionary relationships of species in taxonomically complex groups provide an opportunity to study the earliest stages of speciation, yet few studies to date have investigated such taxa with genomic data. In this study, we applied genotyping-by-sequencing (GBS) to extensive population sampling of British *Euphrasia* species and hybrids across Britain and Ireland. We focused on identifying whether any currently defined species are genomically distinct across their range, and investigated how different evolutionary processes might interact to shape genomic variation and maintain species boundaries. We find that northern *E. micrantha* is genetically cohesive, indicating strong differentiation between this and other sympatric eyebrights, especially within the conserved sub-genome that is shared between diploid and tetraploid taxa. In contrast, the other samples largely show broad-scale genetic clustering corresponding to geography regardless of species identity. Our general finding of genetic clustering by geography rather than species across Britain and Ireland highlights the close evolutionary relationship of species and suggests extensive homogenising gene flow and a key role of local geographic population substructure. We discuss how these findings contribute to our understanding of the origin and maintenance of species in taxonomically complex groups.

### Ploidy as a barrier between species groups

Polyploidy is a major driver of speciation (Alix et al., 2017; Ren et al., 2018); previous studies in *Euphrasia* have shown clustering by ploidy using AFLP markers, suggesting ploidy is a major barrier to gene flow (French et al., 2008). However, an increasing number of studies have shown polyploidy in itself might not be a complete barrier to gene flow (Brown et al., 2022; Husband et al., 2016; Kennedy et al., 2006). Our analyses of all population samples show diploids and tetraploids cluster together at low k-values in fastStructure and that there is little genetic variance explained by ploidy, supporting their close sequence similarity in this shared sub-genome. The genomic similarity between diploids and tetraploids in this sub-genome may facilitate homoeologous chromosome pairing and provide the opportunity for cross-ploidy hybridisation, a factor known to contribute to the origin of new hybrid species (Chapman & Abbott, 2010). For example, *E. vigursii* is a diploid with a putative diploid-tetraploid origin, with tetraploid *E. micrantha* and diploid *E. anglica* as its proposed parental progenitors (French et al., 2008; Yeo, 1956; Silverside, 1990). While we did not find explicit evidence for cross-ploidy hybridisation with our limited sampling of *E. vigursii*, we did see multiple cases of putative cross-ploidy hybridisation, such as a population of samples in northern Scotland with diploid morphological characteristics that clustered with local tetraploids. Above all however, we interpret our clustering results for diploids as indicative of their close relationship to tetraploids, with ploidy still a barrier to hybridization (Brown et al., 2022). In terms of the tetraploid species, these cluster together at low k-values and do not separate into previously recognised species groups at higher k-values (French et al., 2008), indicative of a single origin of the polyploids.

### Genome-wide distinctiveness

We found that currently recognised *Euphrasia* species differ in their extent of genetic divergence, and thus represent different stages within the speciation continuum. Specifically, the Northern group of *E. micrantha* shows genome-wide distinctiveness and is consistently recovered across all our analyses of genomic differentiation (Supplemental Figure 4, Supplemental Table 2). This indicates northern *E. micrantha* is at a later stage in the speciation continuum where populations are largely genomically distinct from other co-occurring taxa. This is consistent with the analysis of genetic clustering using whole genome resequencing data in Fair Isle, Shetland (Becher et al., 2020), which showed *E. micrantha* to be distinct from other tetraploids, based on a small number of samples. Our study indicates that it’s not only an island phenomenon driven by strong genetic drift but extends over the northern extent of the species’ range.

The genetic distinctiveness of northern populations of *E. micrantha* could be explained by several non-mutually exclusive explanations. The first is that Northern *E. micrantha* might be of a different origin or source population to other tetraploid *Euphrasia*. It may be that northern and southern *E. micrantha* are from different glacial refugia, and colonised Britain through different routes, and may now show distinct geographic ranges and with partial reproductive isolation in areas of secondary contact. This pattern of recolonisation from multiple refugia is common in many European plant groups and leaves a distinct genetic signature (Huck et al., 2009; Petit et al., 2002; Valtueña et al., 2012). In *Euphrasia*, previous phylogenetic studies using widespread European sampling showed that there were multiple waves of colonisation to Britain (Garrett et al., 2022; Gussarova et al., 2008), while studies of phylogeography in the *Euphrasia minima* complex found persistence in multiple refugia: one from North America where plants subsequently colonised Greenland/Iceland/Norway, and the other from the Russian Ural mountains where populations subsequently spread to Svalbard (Gussarova et al., 2012).

An alternative, but non-mutually exclusive explanation, is that divergence of *E. micrantha* is caused by ecological specialisation, with differentiation maintained by a high selfing rate. *E. micrantha* is a highly selfing species (Stone, 2012) that is more widely distributed in northern regions of Britain and Europe compared to many of the other widely distributed British *Euphrasia* species (Yeo, 1978). Previous studies have shown a tight evolutionary relationship between *E. micrantha* and its potentially favoured host species *Callunas vulgaris* (Denters, 2013; Yeo, 1968), which both occur in acid heathland and have purple flowers. The presumably higher tolerance to cold temperatures, combined with specialisation in acidic soils, may facilitate the rapid evolution of (partial) reproductive isolation from other congeneric taxa (Cao et al., 2022; Yeo, 1968). Many studies have shown that ecological selection drives genomic divergence and speciation (Cerca et al., 2022; Malinsky et al., 2015; Martin et al., 2013; Querns et al., 2022), and the high selfing rate might contribute to the maintenance of divergence by adapting to greater niche breadth (Grant & Kalisz, 2020). While it is likely that selfing plays an important role in adaptive divergence in *Euphrasia*, this is hard to test with GBS data due to the low coverage per site that will affect estimates of inbreeding, heterozygosity, and linkage disequilibrium. Our future work will investigate selfing using high coverage whole genome resequencing data.

The utility of our genomic data for defining taxa is somewhat limited beyond the northern *E. micrantha* group, given that we are unlikely to have sampled species-specific genomic regions (if they are present). Overall, despite the lack of diagnostic genomic signal, we still believe that many *Euphrasia* species studied here are largely phenotypically and genetically distinct and warrant taxonomic recognition. Previous common garden experiments (Brown et al., 2020) found that a number of *Euphrasia* species maintain their phenotypic distinctiveness under standardised growth conditions, showing a genetic basis for different traits defining the species. However, further taxonomic work is required to assess each species, and it would seem likely that some narrowly defined taxa should be ‘lumped’ into broader species. For example, the presence of isolation by distance across all diploid species suggests these may not be discrete entities, and supports a previous, widened definition of these taxa.

### Range-wide genetic clustering by geography

In terms of the other *Euphrasia* species studied here, our genetic clustering analyses either assign most mainland British tetraploids/diploids to a single genetic cluster (at low k-values, Figure 2C), or to regional genetic clusters (at high k-values, Figure 2D, Supplemental Figure 4, Supplemental Table 2). The geographic clustering pattern in *Euphrasia* is probably due to extensive local hybridisation, while the lack of isolation by distance in the tetraploids is probably a consequence of a selfing or mixed-mating system that causes differentiation at the population level and the idiosyncrasies of introgression (Table 2). Here, sympatric species are likely to show extensive genetic similarity due to a lack of reproductive isolation and extensive pollen mediated gene flow. Despite this extensive homogenising gene flow, other genomic studies of hybridising taxa have shown incipient species may have a ‘mosaic’ genome, with a limited number of genomic regions distinct between species (Andrew & Rieseberg, 2013; Feder et al., 2015; Nadeau et al., 2013). Such ‘genomic islands’ may define taxa and be maintained by strong ecological selection. Further exploration of genome-wide differentiation revealed few sites have elevated F_ST_ between species, however this is based on relatively sparse SNP data from GBS, and without sampling presence-absence-variation that may define taxa (Becher et al., 2021; 2022).

## Conclusions

Our study provides evidence for the hypothesis that species boundaries in recently diverging *Euphrasia* are highly permeable, with geographic genetic structure playing a more significant role in genetic clustering than species identity. The selfing heathland specialist *E. micrantha* in Northern Scotland shows genome-wide divergence from all other taxa, likely due to a distinct post-glacial colonisation history and a highly selfing mating system. However, for all other species we see extensive gene flow and the complete absence of species-specific SNPs. Our study adds to our growing understanding of the extent of sequence similarity between closely related allopatric taxa, with a dramatic case where species are more closely related to neighbouring samples than other conspecific individuals. In the future, whole-genome sequencing of *Euphrasia* species at different stages of the speciation continuum (Thomas-Bulle et al., 2022) will help us understand the accumulation of genetic barriers to gene flow and the factors that promote or prevent divergence.

## Acknowledgements

We thank BSBI members and plant collectors for providing *Euphrasia* samples, and Eric Wafula and Claude dePamphilis for help with assembling the diploid reference genome. We also thank Hannes Becher for providing analysis advice, and Richard Ennos provided useful comments on AMOVA analyses. The Royal Botanic Garden Edinburgh acknowledges funding from the Scottish Government’s Rural and Environment Science and Analytical Services Division (RESAS). YD was supported by the School of Biological Sciences, The University of Edinburgh. AT was supported by the Natural Environment Research Council (NERC) grants NE/R010609/1, NE/L011336/1 and NE/N006739/1.

## Data Accessibility Statement

The raw sequence reads are available on the Sequence Read Archive with accession numbers [Given on acceptance], which can be used for SNP calling in Tassel using the key file provided in Supplementary Materials. The sequence alignments from each dataset, and outputs from fastStructure, PCA, DAPC and conStruct are available in Dryad [Accession information given on acceptance]. The scripts are available on github [Accession information given on acceptance]

## Benefit-Sharing Statement

Benefits Generated: We worked closely with the indigenous community that provided the biodiversity resources for the *Euphrasia* sample collections and identifications. The invaluable contributions of all individuals involved in this research are fully acknowledged in the METHODS and ACKNOWLEDGEMENTS sections. Finally, as previously mentioned, all data have been made accessible to the general public through appropriate biological databases, ensuring that the benefits of this study are widely shared.

## Author Contributions

ADT and YD designed the research; CM identified all *Euphrasia* specimens; ADT carried out sequencing; YD and WH analysed the data; YD and ADT wrote the manuscript. PMH contributed to the manuscript from designing to editing major guidelines.

## Notes

### Competing Interest Statement

The authors have declared no competing interest.

## References

Alix, K., Gérard, P. R., Schwarzacher, T., & Heslop-Harrison, J. S. P. (2017). Polyploidy and interspecific hybridization: partners for adaptation, speciation and evolution in plants. Annals of Botany, 120(2), 183. https://doi.org/10.1093/AOB/MCX079

Andrew, R. L., & Rieseberg, L. H. (2013). Divergence is focused on few genomic regions early in speciation: incipient speciation of sunflower ecotypes. Evolution; International Journal of Organic Evolution, 67(9), 2468–2482. https://doi.org/10.1111/EVO.12106

Andrew, R., & Rieseberg, L. (2013). Divergence is focused on few genomic regions early in speciation: Incipient speciation of sunflower ecotypes. Evolution, 67(9), 2468–2482. https://doi.org/10.1111/evo.12106

Becher, H., Brown, M. R., Powell, G., Metherell, C., Riddiford, N. J., & Twyford, A. D. (2020). Maintenance of species differences in closely related tetraploid parasitic *Euphrasia* (Orobanchaceae) on an isolated island. Plant Communications, 1(6), 100105. https://doi.org/10.1016/J.XPLC.2020.100105

Becher, H., Powell, R. F., Brown, M. R., Metherell, C., Pellicer, J., Leitch, I. J., & Twyford, A. D. (2021). The nature of intraspecific and interspecific genome size variation in taxonomically complex eyebrights. Annals of Botany, 128, 639–651. https://doi.org/10.1093/aob/mcab102

Becher, H., Sampson, J., & Twyford, A. D. (2022). Measuring the invisible: The sequences causal of genome size differences in eyebrights *(Euphrasia)* revealed by k-mers. Frontiers in Plant Science, 13. https://doi.org/10.3389/fpls.2022.818410

Bradburd, G. S., Coop, G. M., & Ralph, P. L. (2018). Inferring continuous and discrete population genetic structure across space. Genetics, 210(1), 33–52. https://doi.org/10.1534/GENETICS.118.301333

Bradburd, G. S. (2019) conStruct: Models spatially continuous and discrete population genetic structure. R pacakges, https://github.com/gbradburd/conStruct

Brown, M. R., Frachon, N., Wong, E. L. Y., Metherell, C., & Twyford, A. D. (2020). Life history evolution, species differences, and phenotypic plasticity in hemiparasitic eyebrights (Euphrasia). American Journal of Botany, 107(3), 456–465. https://doi.org/10.1002/AJB2.1445

Brown, M. R., Becher, H., Williams, S., & Twyford, A. D. (2023). Is there hybridization between diploid and tetraploid Euphrasia in a secondary contact zone? American journal of botany, 110(1), e16100. https://doi.org/10.1002/ajb2.16100

Cao, J., Li, Y., Chang, C., Chung, J. Der, & Hwang, S. Y. (2022). Adaptive divergence without distinct species relationships indicate early stage ecological speciation in species of the *Rhododendron pseudochrysanthum* complex endemic to Taiwan. Plants (Basel, Switzerland), 11(9). https://doi.org/10.3390/PLANTS11091226

Cerca, J., Petersen, B., Lazaro-Guevara, J. M., Rivera-Colón, A., Birkeland, S., Vizueta, J., … Martin, M. D. (2022). The genomic basis of the plant island syndrome in Darwin’s giant daisies. Nature Communications 2022 13:1, 13(1), 1–13. https://doi.org/10.1038/s41467-022-31280-w

Chang, C. C., Chow, C. C., Tellier, L. C. A. M., Vattikuti, S., Purcell, S. M., & Lee, J. J. (2015). Second-generation PLINK: Rising to the challenge of larger and richer datasets. GigaScience, 4(1), 7. https://doi.org/10.1186/S13742-015-0047-8/2707533

Chapman, M. A., & Abbott, R. J. (2010). Introgression of fitness genes across a ploidy barrier. New Phytologist, 186(1), 63–71. https://doi.org/10.1111/J.1469-8137.2009.03091.X

Choi, J. Y., Purugganan, M., & Stacy, E. A. (2020). Divergent selection and primary gene flow shape incipient speciation of a riparian tree on Hawaii Island. Molecular Biology and Evolution, 37(3), 695–710. https://doi.org/10.1093/MOLBEV/MSZ259

Coyne, J. A., & Orr, H. A. (2004). Speciation (illustrate). Oxford University Press.

Denters, T. (2013). *Euphrasia micrantha* Rchb. (Slanke ogentroost) weer in beeld!? Gorteria Dutch Botanical Archives, 36, 2012–2013.

Elshire, R. J., Glaubitz, J. C., Sun, Q., Poland, J. A., Kawamoto, K., Buckler, E. S., & Mitchell, S. E. (2011). A robust, simple Genotyping-by-Sequencing (GBS) approach for high diversity species. PLOS ONE, 6(5), e19379. https://doi.org/10.1371/JOURNAL.PONE.0019379

Ennos, R. A., French, G. C., & Hollingsworth, P. M. (2005). Conserving taxonomic complexity. Trends in Ecology & Evolution, 20(4), 164–168. https://doi.org/10.1016/J.TREE.2005.01.012

Feder, J. L., Egan, S. P., & Nosil, P. (2012). The genomics of speciation-with-gene-flow. Trends in Genetics, 28(7), 342–350. https://doi.org/10.1016/J.TIG.2012.03.009

French, G. C., Ennos, R. A., Silverside, A. J., & Hollingsworth, P. M. (2004). The relationship between flower size, inbreeding coefficient and inferred selfing rate in British *Euphrasia* species. Heredity 2005 94:1, 94(1), 44–51. https://doi.org/10.1038/sj.hdy.6800553

French, G. C., Hollingsworth, A. P. M., Silverside, A. A. J., & Ennos, A. R. A. (2008). Genetics, taxonomy and the conservation of British *Euphrasia*. Conservation Genetics, 9, 1547–1562. https://doi.org/10.1007/s10592-007-9494-9

Garrett, P., Becher, H., Gussarova, G., dePamphilis, C. W., Ness, R. W., Gopalakrishnan, S., & Twyford, A. D. (2022). Pervasive phylogenomic incongruence underlies evolutionary relationships in Eyebrights (*Euphrasia*, Orobanchaceae). Frontiers in Plant Science, 13, 1439. https://doi.org/10.3389/FPLS.2022.869583/BIBTEX

Glaubitz, J. C., Casstevens, T. M., Lu, F., Harriman, J., Elshire, R. J., Sun, Q., & Buckler, E. S. (2014). TASSEL-GBS: A high capacity Genotyping by Sequencing analysis pipeline. PLOS ONE, 9(2), e90346. https://doi.org/10.1371/JOURNAL.PONE.0090346

González-Pérez, Á. M., & Caujapé-Castells, J. (2022). Gene flow, barriers, speciation and hybridization in *Parolinia* species (Brassicaceae) endemic to Gran Canaria. Botanical Journal of the Linnean Society, 198, 403–416. https://doi.org/10.1093/botlinnean/boab069

Goudet, J., & Jombart, T. (2021). hierfstat: Estimation and tests of hierarchical F-statistics. R packages, https://www.r-project.org

Grant, A. G., & Kalisz, S. (2020). Do selfing species have greater niche breadth? Support from ecological niche modeling. Evolution, 74(1), 73–88. https://doi.org/10.1111/EVO.13870

Gussarova, G., Greve Alsos, I., & Brochmann, C. (2012). Annual plants colonizing the Arctic? Phylogeography and genetic variation in the *Euphrasia minima* complex (Orobanchaceae). Taxon, 61(1), 146–160. https://doi.org/10.1002/tax.611011

Gussarova, G., Popp, M., Vitek, E., & Brochmann, C. (2008). Molecular phylogeny and biogeography of the bipolar *Euphrasia* (Orobanchaceae): Recent radiations in an old genus. Molecular Phylogenetics and Evolution, 48(2), 444–460. https://doi.org/10.1016/J.YMPEV.2008.05.002

Huck, S., Büdel, B., Kadereit, J., & Printzen, C. (2009). Range-wide phylogeography of the European temperate-montane herbaceous plant *Meum athamanticum* Jacq.: evidence for periglacial persistence. Journal of Biogeography, 36(8), 1588–1599. https://doi.org/10.1111/J.1365-2699.2009.02096.X

Husband, B. C., Baldwin, S. J., & Sabara, H. A. (2016). Direct vs. indirect effects of whole-genome duplication on prezygotic isolation in *Chamerion angustifolium:* Implications for rapid speciation. American Journal of Botany, 103(7), 1259–1271. https://doi.org/10.3732/AJB.1600097

Jombart, T., & Ahmed, I. (2011). adegenet 1.3-1: new tools for the analysis of genome-wide SNP data. Bioinformatics, 27(21), 3070–3071. https://doi.org/10.1093/BIOINFORMATICS/BTR521

Jombart, T., & Bateman, A. (2008). adegenet: a R package for the multivariate analysis of genetic markers. Bioinformatics, 24(11), 1403–1405. https://doi.org/10.1093/BIOINFORMATICS/BTN129

Jombart, T., Devillard, S., & Balloux, F. (2010). Discriminant analysis of principal components: A new method for the analysis of genetically structured populations. BMC Genetics, 11(1), 1–15. https://doi.org/10.1186/1471-2156-11-94

Kamvar, Z. N., Brooks, J. C., & Grünwald, N. J. (2015). Novel R tools for analysis of genome-wide population genetic data with emphasis on clonality. Frontiers in Genetics, 6(JUN), 208. https://doi.org/10.3389/fgene.2015.00208

Kamvar, Z. N., Tabima, J. F., & Grunwald, N. J. (2014). Poppr: An R package for genetic analysis of populations with clonal, partially clonal, and/or sexual reproduction. PeerJ, 2014(1), 1–14. https://doi.org/10.7717/peerj.281

Kennedy, B. F., Sabara, H. A., Haydon, D., & Husband, B. C. (2006). Pollinator-mediated assortative mating in mixed ploidy populations of *Chamerion angustifolium* (Onagraceae). Oecologia, 150(3), 398–408. https://doi.org/10.1007/S00442-006-0536-7

Li, H., & Durbin, R. (2009). Fast and accurate short read alignment with Burrows– Wheeler transform. Bioinformatics, 25(14), 1754–1760. https://doi.org/10.1093/BIOINFORMATICS/BTP324

Lowry, D. B., Rockwood, R. C., & Willis, J. H. (2008). Ecological reproductive isolation of coast and inland races of *Mimulus guttatus*. Evolution, 62(9), 2196–2214. https://doi.org/10.1111/J.1558-5646.2008.00457.X

Malinsky, M., Challis, R. J., Tyers, A. M., Schiffels, S., Terai, Y., Ngatunga, B. P., Miska, E. A., Durbin, R., Genner, M. J., & Turner, G. F. (2015). Genomic islands of speciation separate cichlid ecomorphs in an East African crater lake. Science (New York, N.Y.), 350(6267), 1493–1498. https://doi.org/10.1126/science.aac9927

Marková, S., Dufresne, F., Rees, D. J., Černý, M., & Kotlík, P. (2007). Cryptic intercontinental colonization in water fleas *Daphnia pulicaria* inferred from phylogenetic analysis of mitochondrial DNA variation. Molecular Phylogenetics and Evolution, 44(1), 42–52. https://doi.org/10.1016/J.YMPEV.2006.12.025

Marques, D. A., Lucek, K., Meier, J. I., Mwaiko, S., Wagner, C. E., Excoffier, L., & Seehausen, O. (2016). Genomics of rapid incipient speciation in sympatric threespine stickleback. PLoS Genetics, 12(2). https://doi.org/10.1371/JOURNAL.PGEN.1005887

Martin, S. H., Dasmahapatra, K. K., Nadeau, N. J., Salazar, C., Walters, J. R., Simpson, F., … Jiggins, C. D. (2013). Genome-wide evidence for speciation with gene flow in *Heliconius butterflies*. Genome Research, 23(11), 1817–1828. https://doi.org/10.1101/GR.159426.113

Mártonfiová, L. (2015). Hybridization in natural mixed populations of sexual diploid and apomictic triploid dandelions *(Taraxacum* sect. Taraxacum): Why are the diploid sexuals not forced out? Folia Geobotanica, 50(4), 339–348. https://doi.org/10.1007/S12224-015-9231-Y

McKinnon, J. S., & Rundle, H. D. (2002). Speciation in nature: the threespine stickleback model systems. Trends in Ecology & Evolution, 17(10), 480–488. https://doi.org/10.1016/S0169-5347(02)02579-X

Metherell, C., & Fred J. R. (2018). Eyebrights *(Euphrasia)* of the UK and Ireland (Vol. 18). Botanical Society of Britain & Ireland.

Nadeau, N. J., Martin, S. H., Kozak, K. M., Salazar, C., Dasmahapatra, K. K., Davey, J. W., … Jiggins, C. D. (2013). Genome-wide patterns of divergence and gene flow across a butterfly radiation. Molecular Ecology, 22(3), 814–826. https://doi.org/10.1111/J.1365-294X.2012.05730.X

Pante, E., & Simon-Bouhet, B. (2013). marmap: A package for importing, plotting and analyzing bathymetric and topographic data in R. PLOS ONE, 8(9), e73051. https://doi.org/10.1371/JOURNAL.PONE.0073051

Padgham, M., & Sumner, D. M. (2021). geodist: Fast, dependency-free geodesic distance calculations. R packages. https://github.com/hypertidy/geodist

Petit, R. J., Brewer, S., Bordács, S., Burg, K., Cheddadi, R., Coart, E., Cottrell, J., Csaikl, U. M., Dam, B., & Deans, J.D. (2002). Identification of refugia and post-glacial colonisation routes of European white oaks based on chloroplast DNA and fossil pollen evidence. Forest Ecology and Management, 156(1–3), 49–74. https://doi.org/10.1016/S0378-1127(01)00634-X

Purcell, S., Neale, B., Todd-Brown, K., Thomas, L., Ferreira, M. A., Bender, D., Maller, J., Sklar, P., de Bakker, P. I., Daly, M. J., & Sham, P. C. (2007). PLINK: a tool set for whole-genome association and population-based linkage analyses. American journal of human genetics, 81(3), 559–575. https://doi.org/10.1086/519795

Querns, A., Wooliver, R., Vallejo-Marín, M., & Sheth, S. N. (2022). The evolution of thermal performance in native and invasive populations of *Mimulus guttatus*. Evolution letters, 6(2), 136–148. https://doi.org/10.1002/evl3.275

Raj, A., Stephens, M., & Pritchard, J. K. (2014). fastSTRUCTURE: Variational inference of population structure in large SNP data sets. https://doi.org/10.1534/genetics.114.164350

Ren, R., Wang, H., Guo, C., Zhang, N., Zeng, L., Chen, Y., Ma, H., & Qi, J. (2018). Widespread whole genome duplications contribute to genome complexity and species diversity in angiosperms. Molecular plant, 11(3), 414–428. https://doi.org/10.1016/j.molp.2018.01.002

Silverside J. A. (1990). A guide to eyebrights *(Euphrasia)* (ii) the *E. rostkoviana* group. Wild Flower Magazine, 418, 31–34.

Seppey, M., Manni, M., & Zdobnov, E. M. (2019). BUSCO: Assessing genome assembly and annotation completeness. Methods in Molecular Biology, 1962, 227–245. https://doi.org/10.1007/978-1-4939-9173-0_14

Stone, H. (2012). The evolution and conservation of tetraploid Euphrasia L. in Britain. The University of Edinburgh, Edinburgh.

Thomas-Bulle, C., Bertrand, D., Nagarajan, N., Copley, R. R., Corre, E., Hourdez, S., … Jollivet, D. (2022). Genomic patterns of divergence in the early and late steps of speciation of the deep-sea vent thermophilic worms of the genus Alvinella. BMC Ecology and Evolution, 22(1). https://doi.org/10.1186/S12862-022-02057-Y

Twyford, A. D., & Friedman, J. (2015). Adaptive divergence in the monkey flower *Mimulus guttatus* is maintained by a chromosomal inversion. Evolution, 69(6), 1476–1486. https://doi.org/10.1111/evo.12663

Twyford, A. D., Wong, E. L. Y., & Friedman, J. (2020). Multi-level patterns of genetic structure and isolation by distance in the widespread plant *Mimulus guttatus*. Heredity, 125(4), 227–239. https://doi.org/10.1038/s41437-020-0335-7

Valtueña, F. J., Preston, C. D., & Kadereit, J. W. (2012). Phylogeography of a Tertiary relict plant, *Meconopsis cambrica* (Papaveraceae), implies the existence of northern refugia for a temperate herb. Molecular Ecology, 21(6), 1423–1437. https://doi.org/10.1111/J.1365-294X.2012.05473.X

Wickham, H. (2016). Ggplot2: Elegant graphics for data analysis (2nd ed.). Springer International Publishing.

Yeo, P. F. (1956). Hybridisation between diploid and tetraploid species of Euphrasia. Watsonia, 3, 253–269.

Yeo, P. F. (1966). The breeding relationship of some European Euphrasia. Watsonia, 6(4), 216–245.

Yeo, P. F. (1968). The evolutionary significance of the speciation of *Euphrasia* in Europe. Evolution, 22(4), 736–747. https://doi.org/10.1111/J.1558-5646.1968.TB03473.X

Yeo, P. F. (1978). A taxonomic revision of *Euphrasia* in Europe. Botanical Journal of the Linnean Society, 77(4), 223–334. https://doi.org/10.1111/J.1095-8339.1978.TB01401.X

