## Supplemental Information for "Genomic clustering by geography not species in taxonomically complex British and Irish eyebrights (*Euphrasia*)"

***Supplemental Figure legends***

**Supplemental Figure 1.** Geographical regions used for tests of geographic clustering for the closely related and widely distributed tetraploid species *E. arctica*, *E. confusa* and *E. nemorosa*.

**Supplemental Figure 2.** Principal components analysis (PCA) showing PC1 & 3 based on the original ATC dataset (All samples, including diploids and tetraploids, mapped to the Tetraploid genome, where we extracted the Conserved scaffolds) of 27,513 SNPs.

**Supplemental Figure 3.** Sample information in the distinctive group (Northern MIC) from PC1 (original ATC dataset of 27,513 SNPs.)

**Supplemental Figure 4.** Genetic structure inferred using fastStructure, for analyses with K = 2 to 15 based on the unlink-ATC dataset of 5220 SNPs.

**Supplemental Figure 5.** Isolation by distance of diploid *Euphrasia* samples, excluding samples from northern Scotland.

**Supplemental Figure 6.** Spatial and non-spatial structure results of diploid samples (from the DDA datasets).

**Supplemental Figure 7.** Isolation by distance of tetraploid samples, excluding the northern selfing genetic cluster primarily composed of *E. micrantha*.

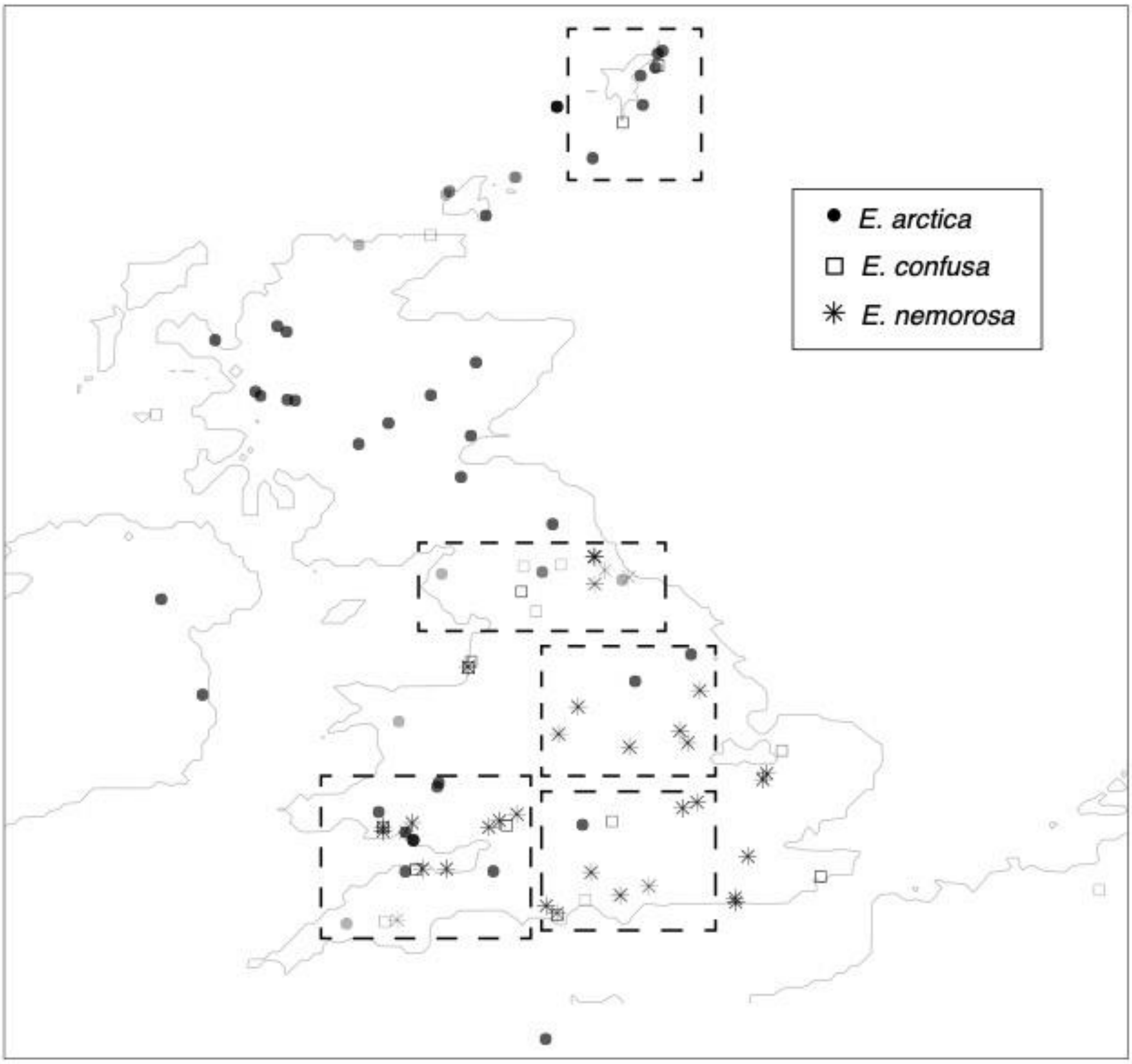


**Supplemental Figure 1.** Geographical regions used for tests of geographic clustering for the closely related and widely distributed tetraploid species *E. arctica*, *E. confusa* and *E. nemorosa*. Regions were arbitrarily defined to 5 areas of similar size, each containing multiple populations from at least two of these three species.

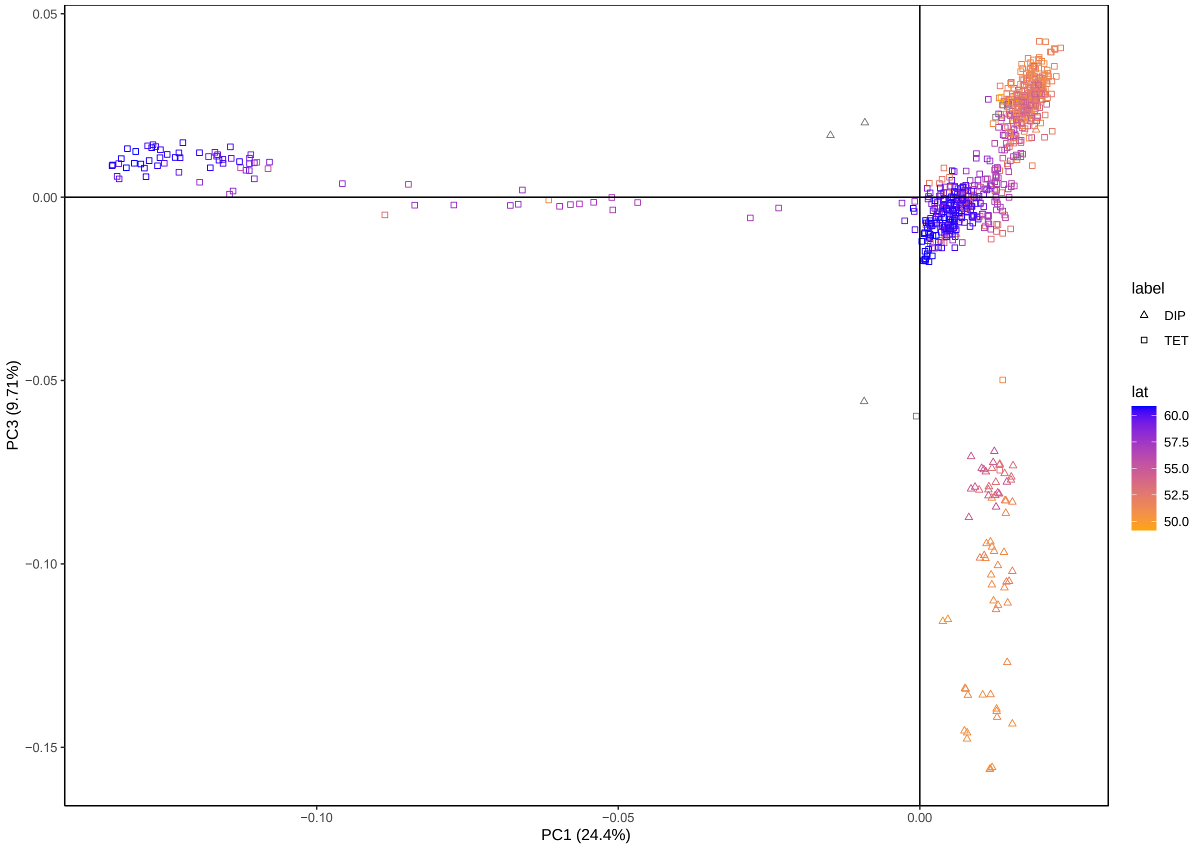


**Supplemental Figure 2.** Principal components analysis (PCA) showing PC1 & 3 based on the original ATC dataset of 27,513 SNPs.

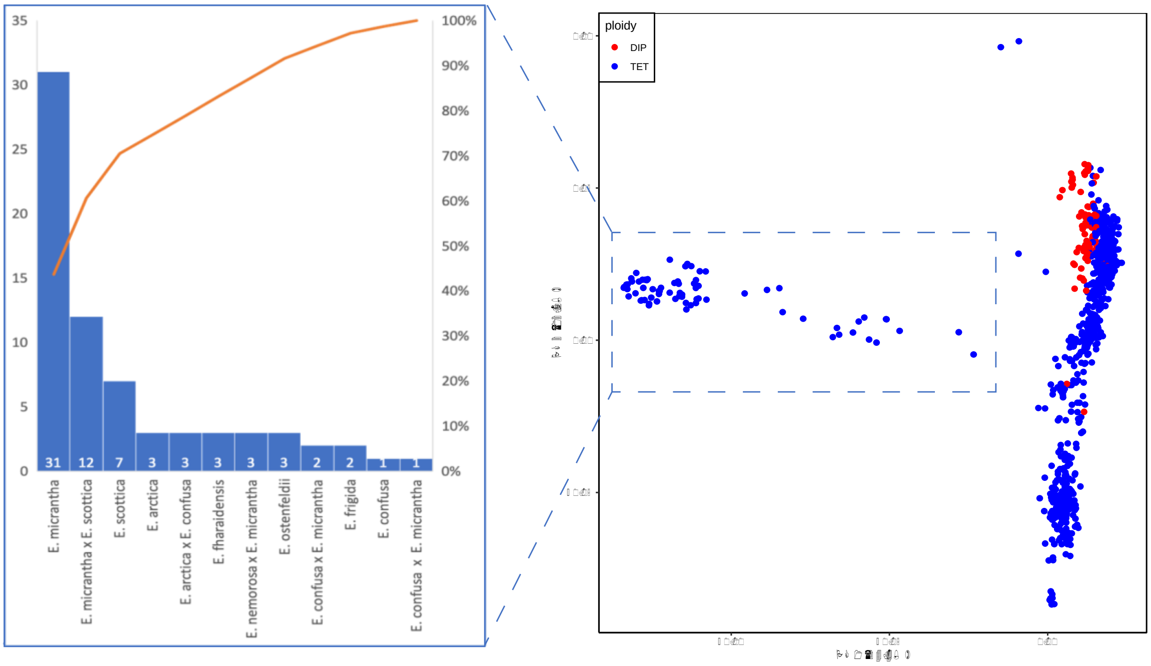


**Supplemental Figure 3.** Sample information in the distinctive group (Northern MIC) from PC1 (original ATC dataset of 27,513 SNPs.)

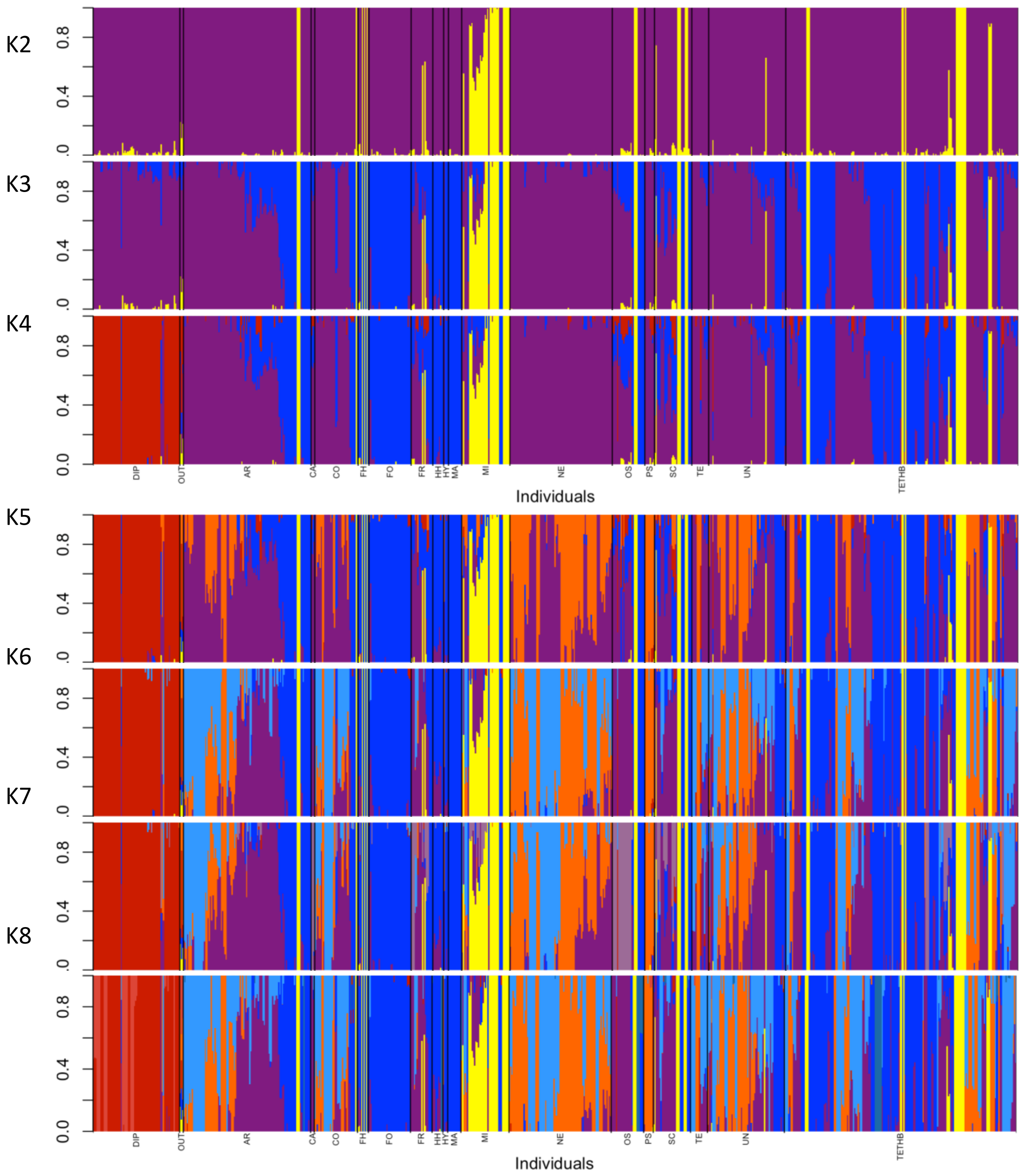

K2
K3
K4
K5
K6
K7
K8
0.0
0.4
0.8
DIP
OUT
AR
CA
CO
FH
FO
FR
HH
HY
MA
MI
NE
OS
PS
SC
TE
UN
TETHB
Individuals

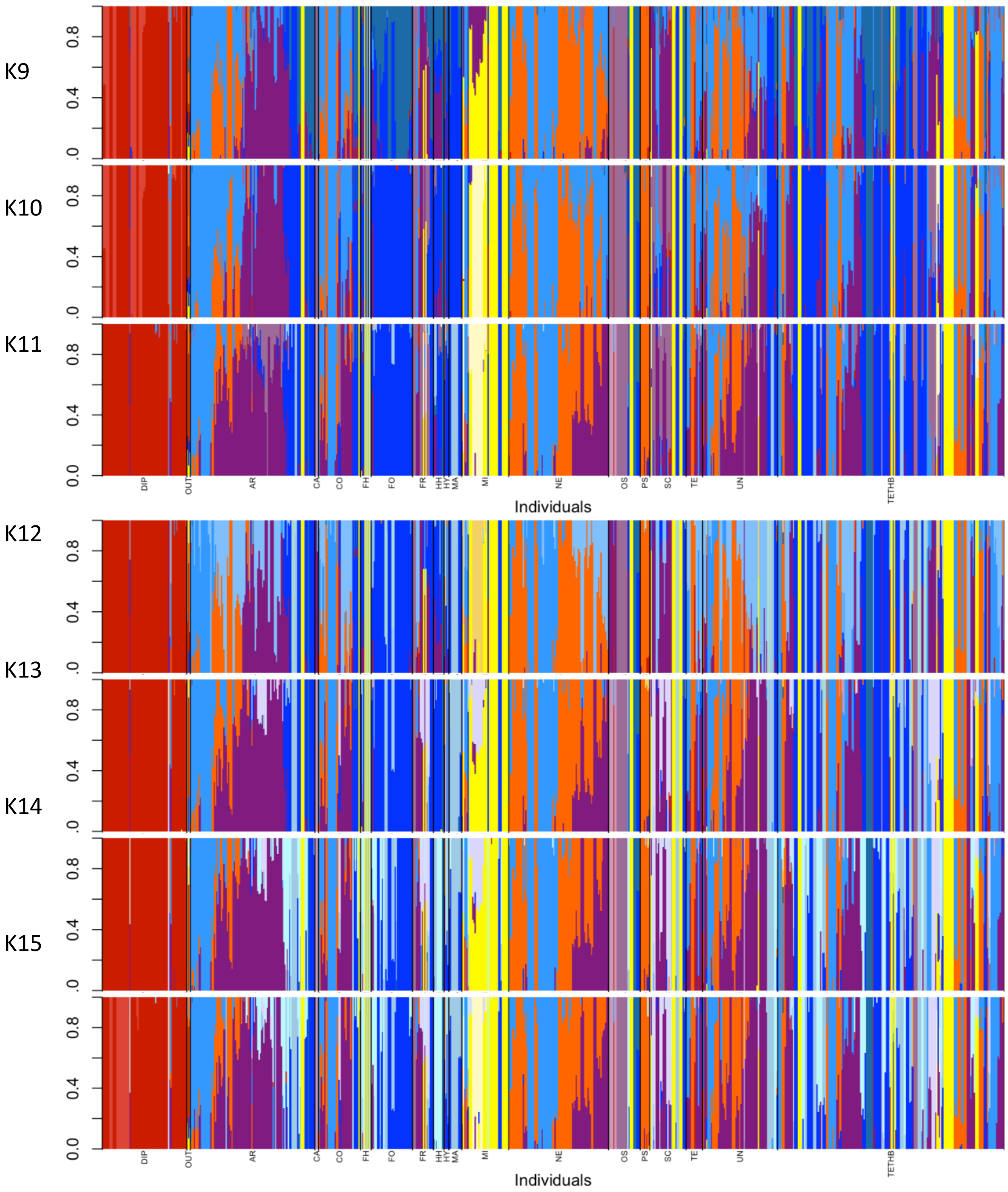


**Supplemental Figure 4.** Genetic structure inferred using fastStructure, for analyses with K = 2 to 15 based on the unlink-ATC dataset of 5220 SNPs. The annotation for each group: DIP-diploid samples; OUT-outgroups (*E. pectinata, E. picta, E. regelli*); AR-*E. arctica*; CA-*E. campbelliae*; CO-*E. confusa*; FH-*E. fharaidensis*; FO-*E. foulaensis*; FR-*E. frigida*; HH-*E. heslop-harrisoniiI*; MA-*E. marshallii*; MI-*E. micrantha*; NE-*E. nemorosa*; OS-*E. ostenfeldii*; PS-*E. pseudokerneri*; SC-*E. scottica*; TE-*E.*

*tetraquetra*; UN-unknown; TETHB-hybrid between tetraploids. New cluster information for analyses from K= 2 to 12 are provided in Supplemental Table 2.

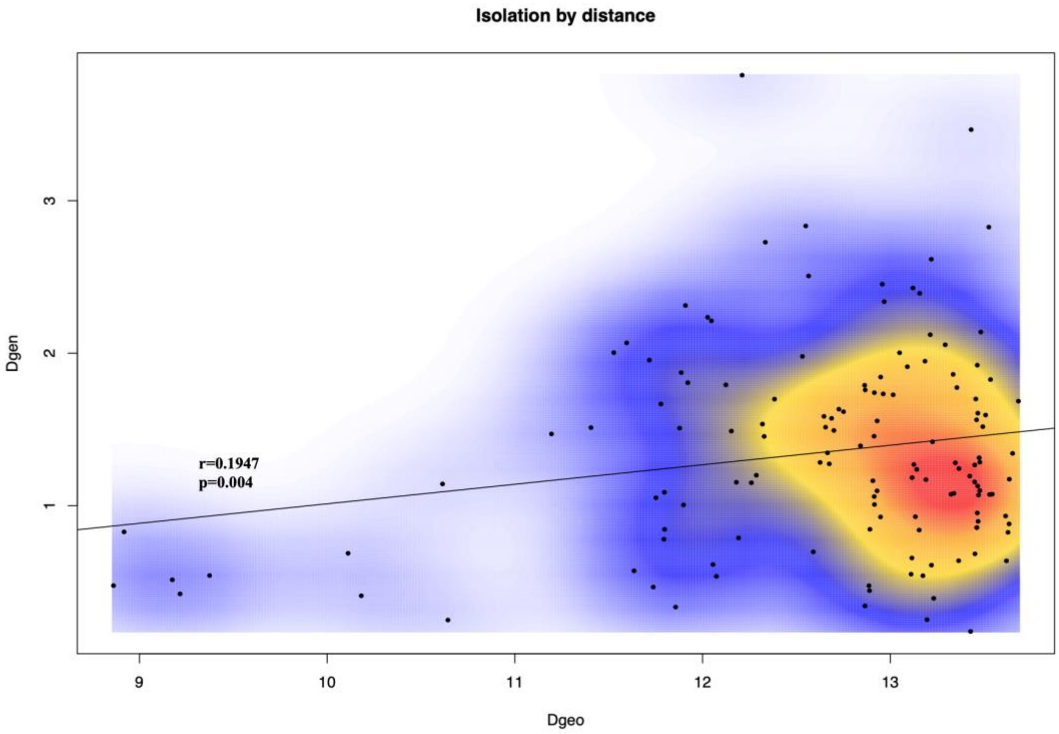


**Supplemental Figure 5.** Isolation by distance of diploid *Euphrasia* samples, excluding samples from vc109 in Scotland.

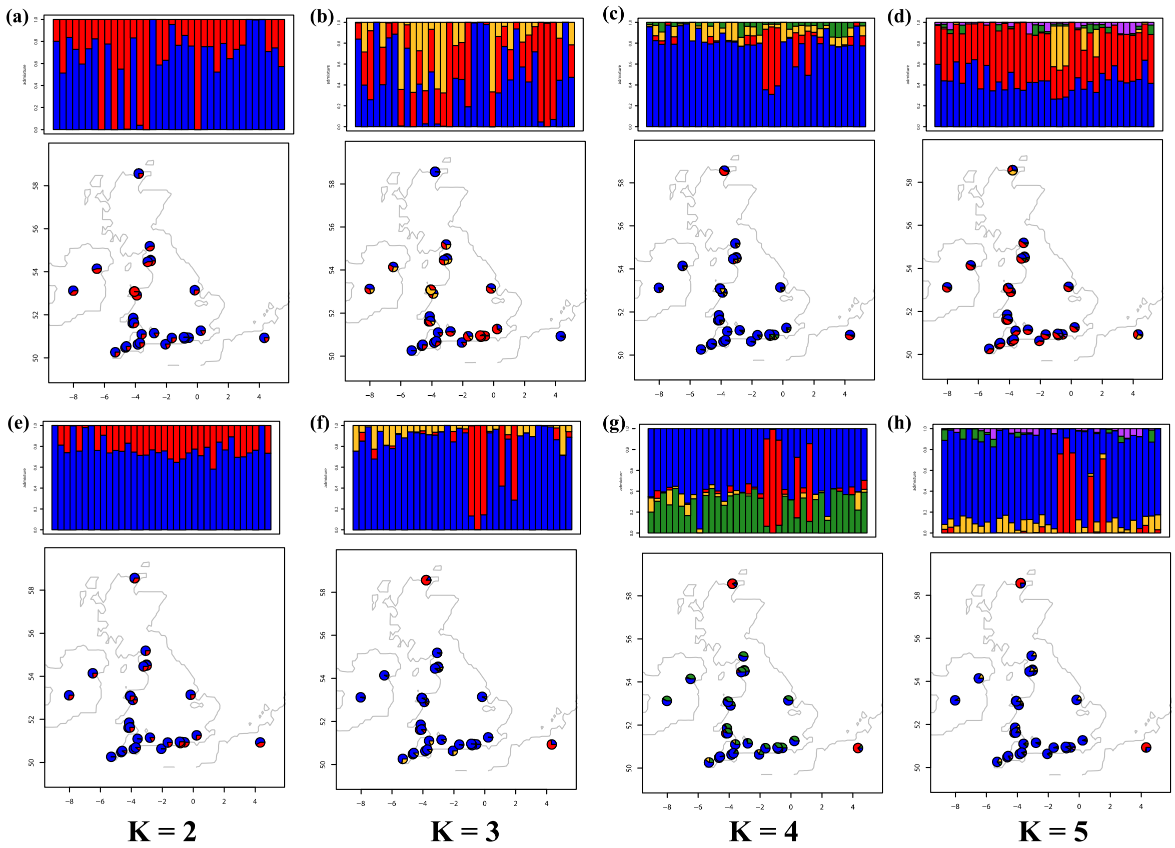


**Supplemental Figure 6.** conStruct results of diploid samples (from the DDA datasets). Top two rows show a nonspatial conStruct model for K=2 through 5(a)-(d) and bottom two rows show spatial construct model for K=2 through 5 (e)-(h).

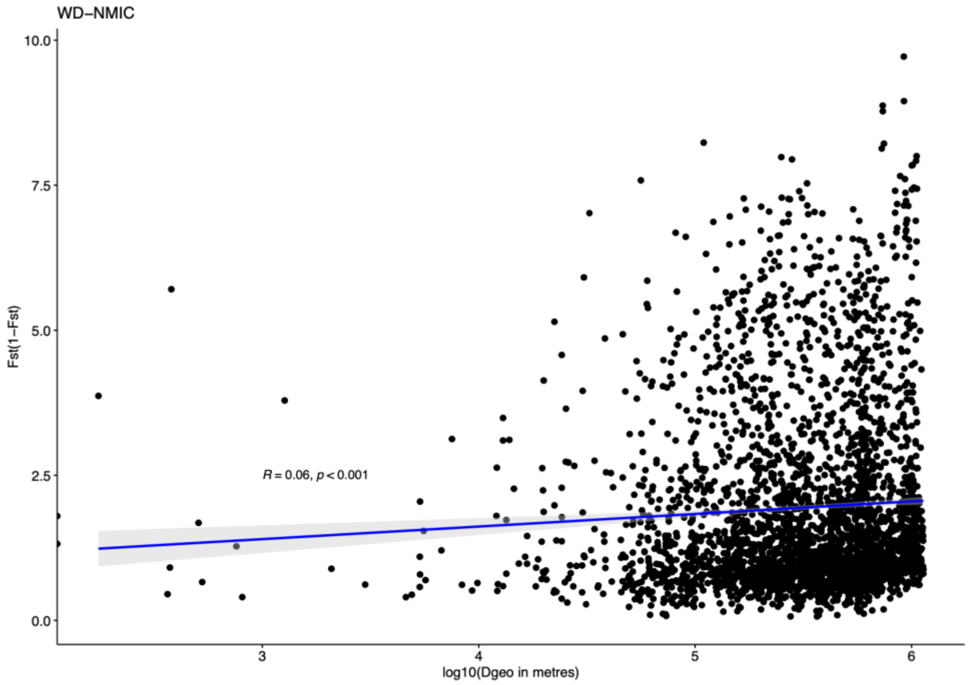


**Supplemental Figure 7.** Isolation by distance of tetraploid samples, excluding the northern selfing genetic cluster primarily composed of *E. micrantha*.

***Supplemental Tables***

**Supplemental Table 1.** Sample information. (Separate file)

**Supplemental Table 2.** New cluster information genetic structure inferred using fastStructure, for analyses with K = 2 to 12 based on the unlink-ATC dataset of 5220 SNPs

| K-value | New genetic cluster |
|---|---|
| 2 | Northern MIC |
| 3 | Geographic split Orkney/Shetland and mainland |
| 4 | Diploids |
| 5 | Southern east group |
| 6 | Southern west group |
| 7 | *E. ostenfeldii* group |
| 8 | Orkney and Shetland |
| 9 | Diploids split into two clusters |
| 10 | Northern Scottish *E. micrantha* came out from Northern *E. micrantha* |
| 11 | *E. marshallii* group |
| 12 | Middle region (roughly) |

**Supplemental Text 1**

Comparisons between the TTC (i.e., tetraploid samples extracted from the ATC dataset, i.e., All samples, including diploids and tetraploids, mapped to the Tetraploid genome, where we extracted the Conserved scaffolds) and TTA (i.e., Tetraploid samples, mapped to the Tetraploid genome,with All tetraploid scaffolds, 10,643 scaffolds) datasets.

SNP called using the TASSEL-GBS pipeline resulted in a total of 70,844 SNPs from the TTA datasets. After sites and individuals were filtered for missing data, there were 16,998 SNPs. The average missing data per site was 19%. After individual filtering, 697 individuals remained for downstream analyses. Filtering unlinked SNPs for the fastStructure analyses resulted in 4266 SNPs for the TTA dataset. A comparison between the results of PCA, DAPC, and fastStructure between TTC and TTA datasets did not show substantial differences.

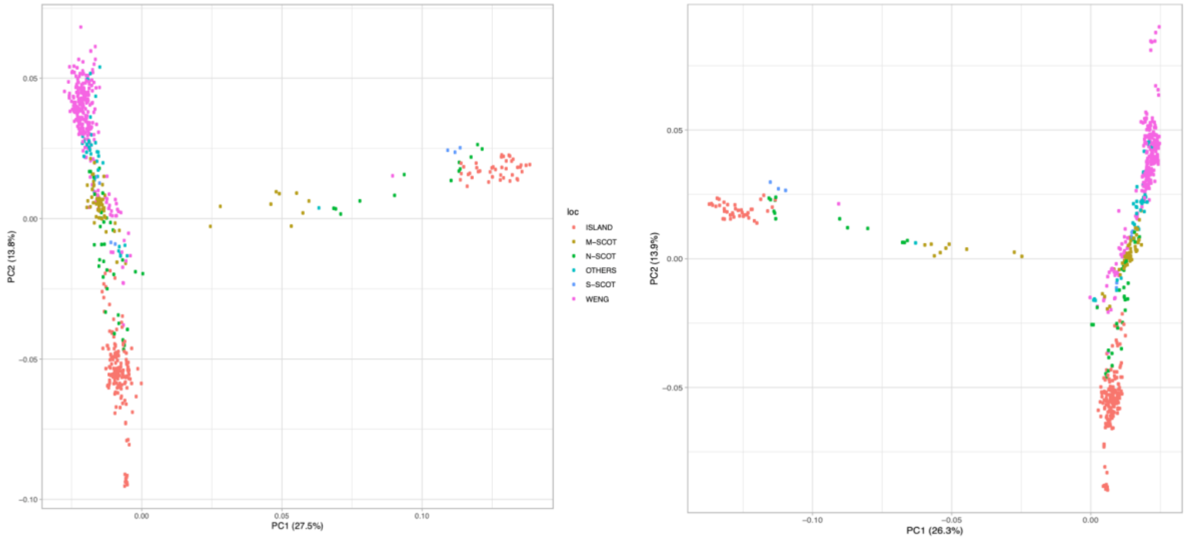


**Supplementary Text 1 Fig.1.** Principal Components analysis between TTC (left) and TTA (right) datasets.

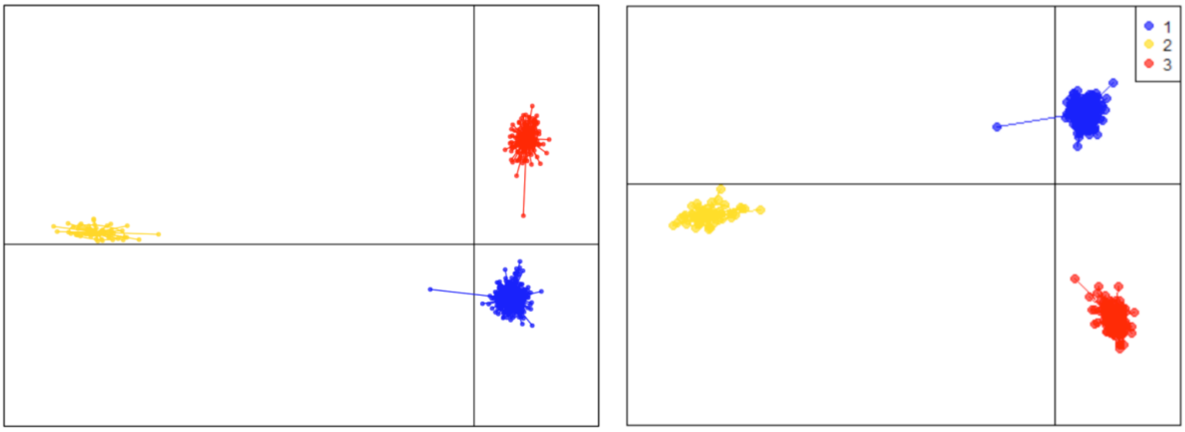


**Supplementary Text 1 Fig.2.** Discriminant analysis of principal components (DAPC) for TTC (left) and TTA (right) datasets.

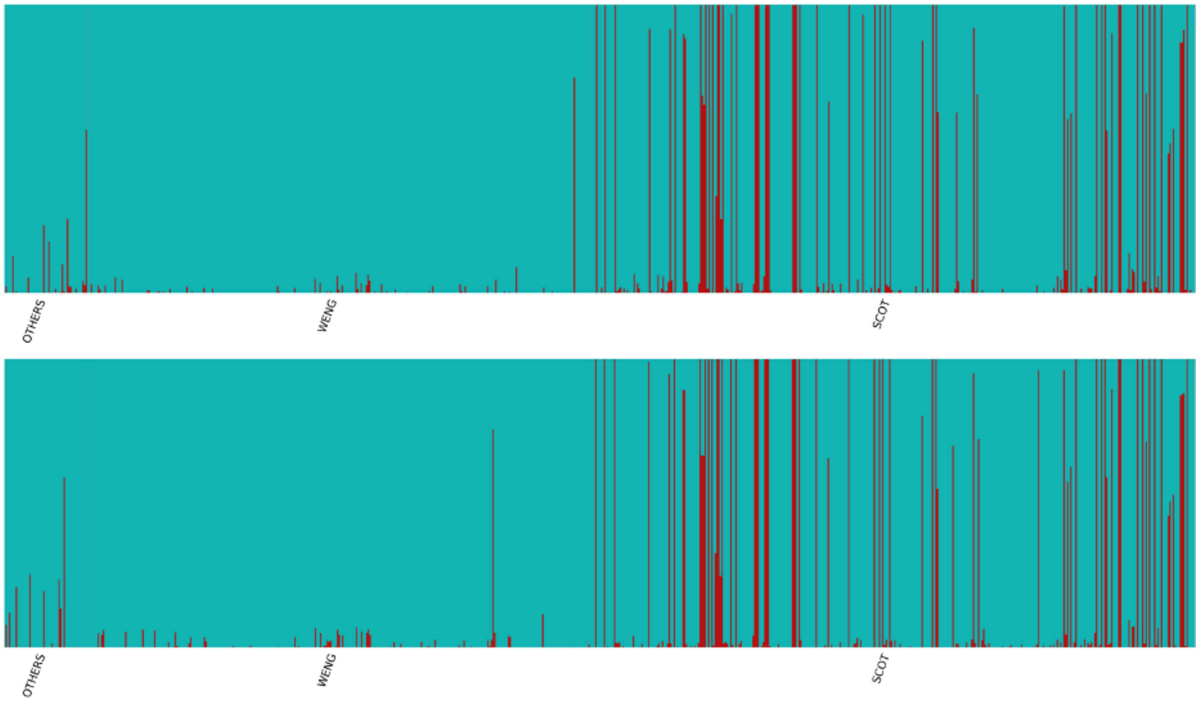


**Supplementary Text 1 Fig.3.** Comparison of results for fastStructure analyses using K=2 between TTC (upper) and TTA (lower) datasets. The annotation in the bottom indicates relative geographic locations. SCOT is Scotland, WENG is Wales and England, OTHERS are all other sites outside Great Britain.

**Supplementary Text 2**

To test whether genetic clusters observed in the population genetic structure analyses were affected by extensive sampling of particular geographic regions or species groups (especially the bias of 800 tetraploid samples vs 80 diploids), we down sampled the ATC dataset to a similar number of individuals between the most distinctive genetic clusters recovered in preliminary analyses, before running for analyses on this large ATC dataset. In this pre-running, we included 50 diploid individuals, 43 *E. micrantha* individuals, and 105 other tetraploid individuals, as widely distributed as it can be, keeping 1 individual per population (detailed individual information in Supplementary Table 1, marked as 'Yes' in 'downsample' column). The results from both PCA and DAPC confirmed the consistency with the larger ATC dataset.

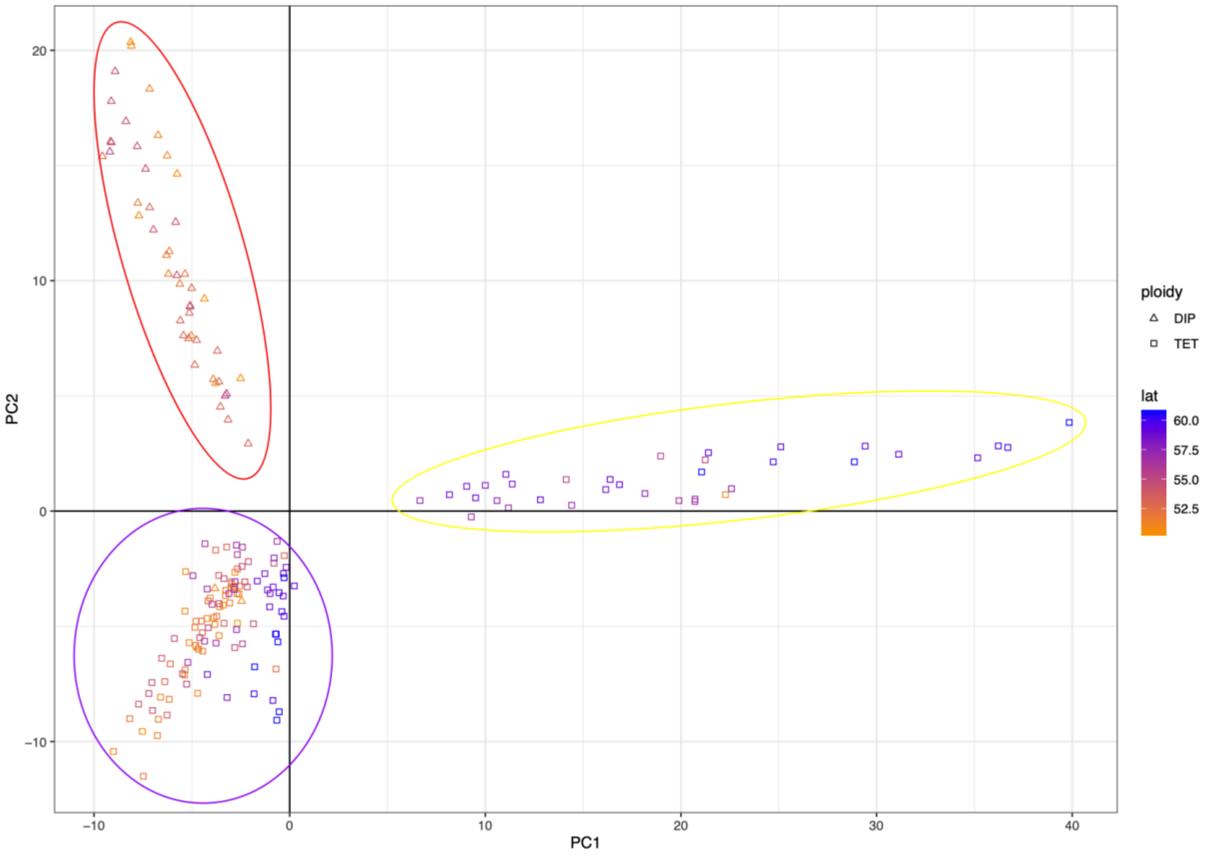


**Supplementary Text 2 Fig.1.** Principal Components analysis (PCA). Yellow circle represents Northern MIC, red circle represents diploid samples, purple circle represents other tetraploid samples.

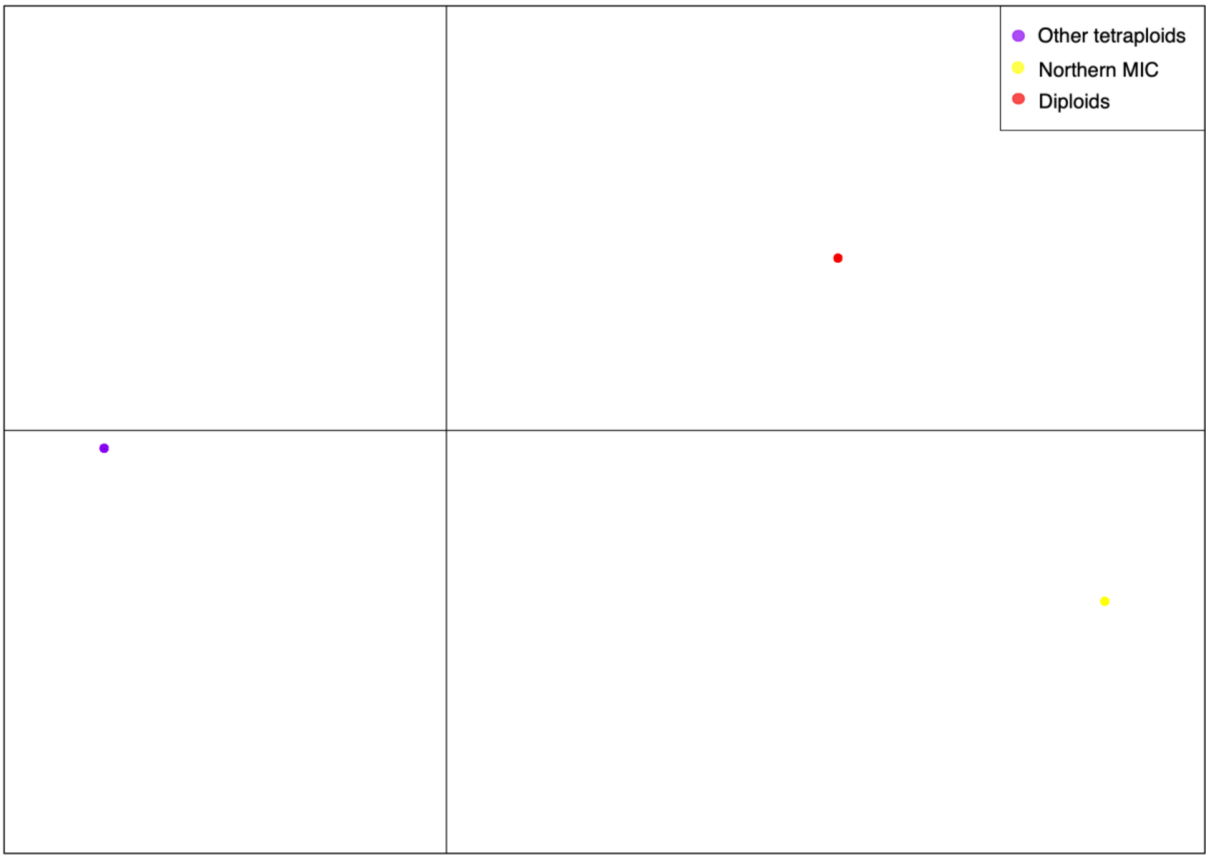


**Supplementary Text 2 Fig.2.** Discriminant analysis of principal components (DAPC).
